# Structural variants and short tandem repeats impact gene expression and splicing in bovine testis tissue

**DOI:** 10.1101/2023.06.07.543773

**Authors:** Meenu Bhati, Xena Marie Mapel, Audald Lloret-Villas, Hubert Pausch

## Abstract

Structural variants (SVs) and short tandem repeats (STRs) are significant sources of genetic variation. However, the impacts of these variants on gene regulation have not been investigated in cattle. Here, we genotyped and characterized 19,408 SVs and 374,821 STRs in 183 bovine genomes and investigated their impact on molecular phenotypes derived from testis transcriptomes. We found that 71% STRs were multiallelic. The vast majority (95%) of STRs and SVs were in intergenic and intronic regions. Only 37% SVs and 40% STRs were in high LD (R2>0.8) with surrounding SNPs/Indels, indicating that SNP-based association testing and genomic prediction are blind to a non-negligible portion of genetic variation. We showed that both SVs and STRs were more than two-fold enriched among expression and splicing QTL (e/sQTL) relative to SNPs/Indels and were often associated with differential expression and splicing of multiple genes. Deletions and duplications had larger impacts on splicing and expression than any other type of structural variant. Exonic duplications predominantly increased gene expression either through alternative splicing or other mechanisms, whereas expression- and splicing-associated STRs primarily resided in intronic regions and exhibited bimodal effects on the molecular phenotypes investigated. Most e/sQTL resided within 100 kb of the affected genes or splicing junctions. We pinpoint candidate causal STRs and SVs associated with the expression of *SLC13A4* and *TTC7B*, and alternative splicing of a lncRNA and *CAPP1*. We provide a catalogue of STRs and SVs for taurine cattle and show that these variants contribute substantially to gene expression and splicing variation.

## Introduction

Genome-wide association studies (GWAS), and expression and splicing quantitative trait loci (e/sQTL) mapping establish links between genotype and (molecular) phenotype (Visscher et al. 2012; Yang et al. 2013; Littlejohn et al. 2016; Lopdell et al. 2017; Sinnott-Armstrong et al. 2021). These approaches typically rely on single nucleotide polymorphism (SNP) and small insertion and deletion (Indel, smaller than 50 bp) markers because they can be genotyped easily and accurately with short sequencing reads using reference-guided approaches. Complex DNA variations such as structural variants (SVs, larger than 50 bp) or short tandem repeats (STRs) are often neglected for GWAS and e/sQTL mapping because they are challenging to genotype. However, it becomes increasingly apparent that SVs and STRs contribute substantially to trait variation (Baker 2012; Sudmant et al. 2015; Abel et al. 2020; Bertolotti et al. 2020; Collins et al. 2020).

Structural variants can be classified into deletions, duplications, insertions, inversions, translocations, segmental duplications, mobile element insertions or complex rearrangements, which may be a combination of multiple types (Ho et al. 2020; Belyeu et al. 2021). Tandem repeats are consecutive repeats of units ranging from 1 bp to several kb (Gymrek 2017). Short tandem repeats (STRs) specifically refer to repeats of a motif between 1 and 6 bp in length, e.g., AGC_7_ indicates that a trinucleotide (3 bp) AGC motif is repeated 7 times, yielding a total length of 21 bp. Polymorphic STRs can vary in length due to a contraction or expansion of the repeat motif. These variants can arise due to recombination errors, insertions of mobile genetic elements, slippage during DNA replication or imperfect DNA repair (Ellegren 2000; Sun et al. 2012; Escaramís et al. 2015).

Historically, structural variants and short tandem repeats had been genotyped using microarray- and polymerase chain reaction (PCR)-based methods. The resulting genotypes were used to validate parentage, construct genetic linkage maps, assess genetic diversity, and map quantitative trait loci (QTL) (Machugh et al. 1994; Ihara et al. 2004; Wang et al. 2007; McClure et al. 2012). However, these methods investigate only a small number of polymorphic SVs and STRs. Exhaustive genome-wide discovery and genotyping of SVs and STRs has become feasible through advancements in short read sequencing and variant detection methods (Willems et al. 2014; Sudmant et al. 2015; Saini et al. 2018; Nelson et al. 2019; Abel et al. 2020; Collins et al. 2020). Yet, there are only few studies that identified SVs using whole-genome sequencing data in cattle (Boussaha et al. 2015; Chen et al. 2017; Mesbah-Uddin et al. 2018; Kommadath et al. 2019; Lee et al. 2023). To the best of our knowledge, STRs have not been profiled systematically in different cattle breeds using whole genome sequencing data, as there is only one study which characterized 60,106 STRs in five Holstein cattle (Xu et al. 2017).

It is well known from investigations in species other than cattle that STRs and SVs contribute substantially to complex traits and diseases through mediating gene expression and splicing (Cao et al. 2020; Scott et al. 2021; Vialle et al. 2022). For instance, an intronic AAGGG expansion in the *RFC1* gene encoding Replication Factor C1 is associated with cerebellar ataxia with neuropathy and bilateral vestibular areflexia syndrome in humans (Rafehi et al. 2019). Analyses of the human Genotype-Tissue Expression (GTEx) data showed that SVs were the lead variants in 2.66% cis-eQTL (Scott et al. 2021) and revealed many STRs affecting gene expression (Fotsing et al. 2019). A recent study by Hamanaka et al. (2023) showed that tandemly repeated motifs of up to 20 bp contribute substantially to alternative splicing and thereby phenotype variation (Hamanaka et al. 2023). Recent efforts from the cattle genotype-tissue expression (GTEx) project (Liu et al. 2022) have revealed non-coding SNPs and Indels that impact splicing and expression variation in multiple tissues. However, they did not explore the roles of SVs and STRs. To date, the contribution of SVs and STRs to gene expression and splicing variation are largely unknown in cattle. Therefore, we generated a catalogue of polymorphic STRs and SVs from 183 whole-genome sequenced cattle and assessed the impact of these variant types on gene expression and splicing in testis transcriptomes of 75 mature bulls. Finally, we pinpoint candidate causal STRs and SVs that modulate the expression and splicing of genes in testis tissue.

## Material and Methods

### Alignment and variant calling (SNPs & Indels)

We used paired-end (2 x 150 bp) whole-genome sequencing data of 183 individual cattle (mean coverage 12x) from the Brown Swiss (BSW), Original Braunvieh (OB), Grauvieh (GV), Holstein (HOL) and Fleckvieh (FV) breeds, and their crosses. Reference-guided alignment and variant discovery were performed as described in (Lloret-Villas et al. 2021). Breed information and coverage for all animals, along with their accession numbers, can be found in Table S1 in File S2. In brief, we aligned reads that passed quality control to the ARS-UCD1.2 reference genome using the MEM-algorithm of the Burrows-Wheeler Alignment (BWA) software (Li 2013) with option -M. Read duplicates were marked with the MarkDuplicates module from the Picard Tools software suite (Broad Institute. Picard tools (2021). Subsequent discovery and genotyping of SNPs and Indels was performed with GATK HaplotypeCaller (version 4.1) (Depristo et al. 2011). We filtered the variants with hard filtration settings recommended by GATK to retain high-quality SNPs and Indels. Finally, we imputed sporadically missing genotypes with Beagle (version 4.1) (Browning and Browning 2016) and retained variants with minor allele frequency > 0.05 for downstream analysis.

### Building reference STRs

A previously proposed HipSTR (Willems et al. 2017) workflow (https://github.com/HipSTR-Tool/HipSTR-references/tree/master/mouse) was applied to compile a set of reference STRs from the soft-masked ARS-UCD1.2 reference genome (available from Ensembl (v. 104)). Briefly, we ran the Tandem Repeat Finder (TRF) software for each chromosome with settings 2,7,7,80,10,5,500 -h -d -l 6 -ngs (Benson 1999). We retained repeats with a motif size between 1 and 6 bp, merged overlapping repeats, and finally kept sites with high scores according to motif size as implemented in the trf_parser.py utility. STRs that are not within 10 bp from another STR were retained.

### STRs genotyping

The STRs were genotyped in the cohort of 183 cattle using the default mode of the HipSTR software tool (Willems et al. 2017). The resulting VCF file was filtered using the filter_vcf.py script from HipSTR, with options --min-call-qual 0.8, --max-call-flank-indel 0.20 and --min-loc-depth 5x. We kept only STRs with genotyping rate higher than 60% and at least 1 bp difference. Furthermore, the minimum STR length was established at 11 base pairs, which means that mononucleotide STR required a minimum of 11 copies.

### SVs calling

We applied the smoove workflow to discover and genotype SVs from short sequencing reads (Pedersen et al. 2020). This approach extracts split and discordant reads from each bam file using samblaster (Faust and Hall 2014). These reads are then further filtered using lumpy (Layer et al. 2014) based on several quality metrics. The filtered reads were subsequently used to genotype SVs in each sample separately. The sample-specific SV calls were merged to obtain a set of SVs that segregate in the cohort. Each sample was then re-genotyped for the common set of SVs using SVTyper (Chiang et al. 2015), and Duphold (Pedersen and Quinlan 2019) was run to add depth fold-change. A single joint VCF file was eventually generated that contained deletions (DEL), duplications (DUP), inversions (INV) and breakends (BND). We retained only SVs that were longer than 50 bp, for which the breakpoints were precisely known, and that were supported by at least 1 split read. We kept DUP based on average DHFFC (fold-change of the variant depth relative to flanking regions) scores as het > 1.25 and homo alt > 1.3 and DEL with DHFFC het < 0.70 and DHFFC homo < 0.50. INV were kept if their quality score was above 100. If multiple SVs were reported for the same location, we kept the variant with the highest quality score.

### Annotation of variants

Both STRs and SVs were annotated according to the Ensembl annotation (v. 104) of the bovine genome in a hierarchical manner using BEDTOOLS intersect (Quinlan and Hall 2010) (exon > intron > promoter > intergenic). The promoter was defined as the region located within 5 kb upstream of the gene start (5’ end). We assessed if exonic SVs overlap the whole gene or if they overlap only partially as proposed by Collins et al. (Collins et al. 2020). SNPs and Indels were annotated with the Variant Effect Predictor (VEP) tool (McLaren et al. 2016) based on the Ensembl annotation of the bovine genome (version 104). The most severe consequence for each variant was then assigned to exon, intron, promoter and intergenic regions as above.

### Population structure and linkage disequilibrium (LD)

We used Plink1.9 (Purcell et al. 2007) to calculate the principal components of genomic relationship matrices constructed from SNPs/Indels, SVs or STRs. We used Bcftools (Danecek et al. 2021) to extract all SNPs and Indels within 50 kb of SVs or STRs. For each SV and STR, we calculated LD as the squared Pearson correlation coefficient (R^2^) with the dosage of each surrounding SNP or Indel (maf > 0.05) where dosage is 0 for the 0/0, 1 for the 0/1 and 2 for the 1/1 genotype (Fotsing et al. 2019).

### Pre-processing RNA seq data and alignment

Total RNA of testis tissue from 76 mature bulls that are a subset of the 183 bulls used to profile STRs and SVs were available from a previous study (Kadri et al. 2021). The stranded paired-end reads were trimmed for adapter sequences, low quality bases, and poly-A and poly-G tails with fastp (Chen et al. 2018). The filtered reads were aligned to the ARS-UCD1.2 reference genome and the Ensembl gene annotation (v.104) using STAR (version 2.7.9a) with options - - twopassMode, --waspOutputMode, and --varVCFfile (Dobin et al. 2013).

### Gene expression quantification

Gene level expression (in transcript per million (TPM)) was quantified with the QTLtools quan function with default settings (Delaneau et al. 2017). Raw read counts were obtained with FeatureCounts (Liao et al. 2014). We retained genes that had expression values >0.2 TPM in at least 20% of samples and > 6 reads in at least 20% of samples. A PCA was conducted using log_2_(TPM +1) transformed expression values. One sample was excluded as it appeared as an outlier in the PCA. Finally, TPM values were quantile normalized and inverse normal transformed across samples per gene using the R package RNOmni (McCaw et al. 2020).

### Splicing quantification

We used RegTools (Cotto et al. 2023) and LeafCutter (Li et al. 2018) to quantify intron excision ratios. First, we filtered the STAR-aligned bam files for uniquely aligned and wasp-filtered reads (tag vW:i:1) (van de Geijn et al. 2015). Next, exon-exon junctions were obtained using RegTools with option -a 8 -m 50 -M 500000 -s 1. Finally, introns were clustered with a modified version of the leafcutter_cluster.py script provided by the Human GTEx consortium (The GTEx Consortium 2020). The script additionally filters introns without any read counts in >50% of samples and insufficient variability. Finally, the filtered intron counts were normalized using the prepare_phenotype_table.py script from LeafCutter and converted to BED format with the start/end position corresponding to the first position of 5’ of intron cluster.

### Covariates for e/sQTL analysis

To account for hidden confounders that might cause variance of gene expression or splicing, we applied the Probabilistic Estimation of Expression Residuals (PEER) (Stegle et al. 2012). The top three principal components of a genomic relationship matrix that was calculated based on LD pruned (--indep-pairwise 50 10 0.1) SNPs using Plink1.9 (Purcell et al. 2007) were used to account for population structure. The influence of covariates on gene expression and splicing was quantified with the variancePartition R package (Hoffman and Schadt 2016).

### e/sQTL mapping

We used the difference in length between reference and alternate (computed from the sum of the GB format tag in the output VCF file from HipSTR) alleles as dosage for the STRs for eQTL mapping (Fotsing et al. 2019; Jakubosky et al. 2020). To minimize the effect of outlier STRs, we converted the genotypes to missing if they were not observed in at least two samples. We kept sites with >80% genotyping rate. To prevent the removal of multiallelic sites by QTLtools, we replaced the alternate allele field of the VCF file with the string “STR”. Furthermore, the GT field (genotype) was substituted with dosage values. Genotypes of SVs, SNPs and Indels, were also converted to dosages (0/0 to 0, 0/1 to 1 and 1/1 to 2). Genotypes at each variant position were normalized so that the effect size can be compared across the different variant types. All these changes were implemented using custom Python scripts. We performed cis-eQTL mapping between expressed genes and all variants in *cis* (± 1 Mb) with QTLtools using the cis permutation mode (1000 permutations) and accounting for covariates (5 PEER factors, 3 PC, RIN and age). To account for multiple testing per molecular phenotype (Genes), we used the Storey & Tibshirani False Discovery Rate procedure that was implemented with the R/qvalue package on beta approximated p-values (eGene) as described by (Delaneau et al. 2017). This approach resulted in genes (eGenes) that had at least one significant eVariant and threshold p values for all genes. Finally, we performed conditional analyses using QTLtools with threshold p values to identify all significant independent eVariants per eGene which were used for all subsequent comparison.

Cis-sQTL mapping was performed as described above with QTLtools using the cis permutation mode and accounting for covariates (5 PEER factors, 3 PC, RIN and age). We employed grouped permutations (--grp_best option) to collectively calculate an empirical p-value across all introns within an intron cluster. Normalized intron excision ratios (the ratio of the reads defining an excised intron to the total number of reads of an intron cluster) were used as molecular phenotypes. We considered sQTL to be an sVariant per sIntron cluster pair. Significant intron clusters were annotated (candidate intron boundaries per cluster) based on the ARS-UCD1.2 gene annotation and strand match (Ensembl release 104). Intron cluster coordinates that mapped to multiple genes were considered as unannotated although the number of such intron clusters was less than 100 in each sQTL analyses.

### Properties of e/s Variants

From each sQTL/eQTL analyses, we annotated each e/sVariant type with their respective annotation category as described above. For all enrichment analyses, we used Fisher’s Exact Test (two sided). All plots were created in R (v 3.6.3) with ggplot2. Gene structure was plotted with the ggtranscript R package (Gustavsson et al. 2022) and plots were combined with patchwork (Pedersen 2023).

## Results

We used paired-end whole-genome sequencing data of 183 cattle from five breeds to genotype SVs, STRs, SNPs and Indels. The average sequencing coverage was 12.8-fold and it ranged from 5.0 to 30.4-fold.

### Reference-guided discovery of short tandem repeats

We identified 1,202,536 STRs with a motif size between 1 and 6 bp in the current *Bos taurus taurus* reference sequence (ARS-UCD1.2) spanning 24.9 Mb autosomal sequence (1.0%) (Figure S1 in File S1 and Table S2 in File S2). The number of STRs on each chromosome was correlated (r=0.99) with chromosome length (Figure S2 in File S1). Mono- and hexanucleotide loci were the most and least frequent types of STRs respectively, amounting to 35.9% and 9.8% of all identified STRs (Figure S1 in File S1). Repeats of A, T and AT were most prevalent among mono- and dinucleotide STRs. GC-rich repeats (e.g., AGC) were most frequent among trinucleotide STRs (Figure S3 in File S1). The overall length of the STRs varied from 11 bp to 10,427 bp with a median size of 18 bp. The vast majority of the STRs (n=1,199,357, 99.7%) were shorter than 100 bp, facilitating short read-based genotyping.

### Genotyping of short tandem repeats

We obtained genotypes for 794,300 autosomal STRs in 183 cattle using HipSTR (Willems et al. 2017), of which we retained 374,822 polymorphic loci after stringent filtering for downstream analyses (Table S3in File S2). We identified between 73,791 and 189,658 (average: 150,104) STRs in each cattle genome, and the number of STRs detected correlated (r=0.94) with sequencing depth (Figure S4 in File S1). As expected, given their prevalence in the bovine reference genome, mono- (52.9%) and hexanucleotide STRs (2.5%) were respectively the most and least frequent type of the polymorphic STRs (Figure 1a). Pentanucleotide STRs were more frequent than tetranucleotide STRs. Approximately three quarter of polymorphic STRs (n=266,509, 71.1%) were multiallelic and had between 1 and 41 alternate alleles, but more than 20 alternate alleles were rarely seen (Figure S5 in File S1). Dinucleotide STRs had the highest number of alternative alleles among all STRs (Figure 1b). Repeats of A and T were the most frequent classes among the mononucleotide STRs, whereas AGC and CTG repeats prevailed among trinucleotide STRs (Figure S6 in File S1). Heterozygosity and allelic diversity were higher for dinucleotide loci than any other type of STRs (Figure 1b & c), possibly suggesting higher mutation rate and less purifying selection in this class.

**Figure 1:**
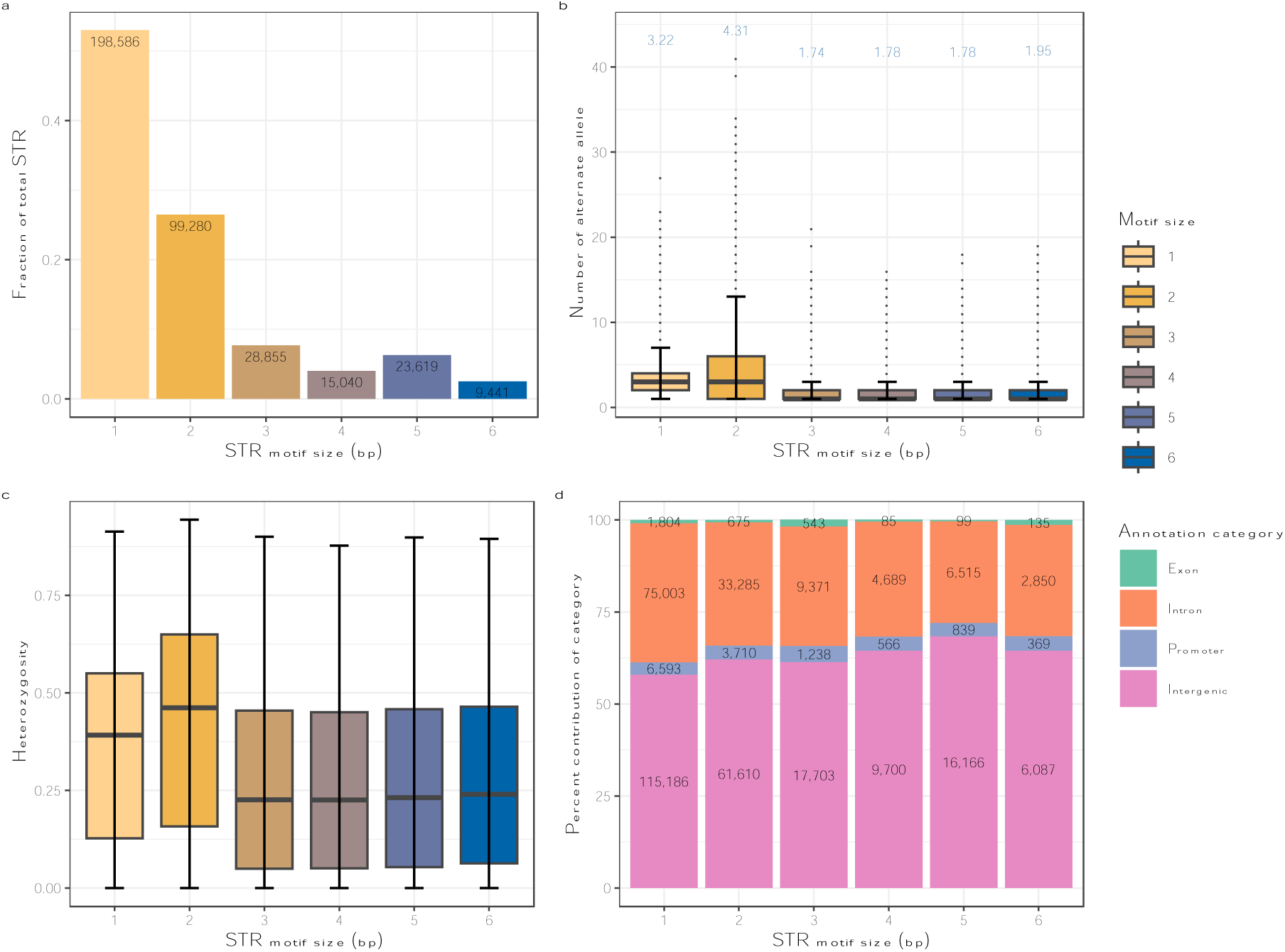
Properties of 374,822 polymorphic STRs in 183 taurine *Bos taurus taurus* cattle genomes. (a) Proportion and count of STRs for each motif size. (b) Number of alternative alleles observed for each STR motif size. Numbers above the boxplots indicate the average number of alleles observed in the 183 cattle genomes for each motif size. (c) Heterozygosity in each STRs motif. (d) Proportion of loci overlapping four annotation categories for each STR motif size. Numbers inside the stacked bars represent the total count of STRs for each annotation category.

We investigated the overlap between the STRs detected in our study and 16 STRs that had been frequently used for parentage testing (including those that had been approved by the International Society for Animal Genetics (ISAG)) (Van De Goor et al. 2009) (Table S4 in File S2). Forward and reverse primer sequences of the 16 STRs were subjected to a BLASTn search against the ARS-UCD1.2 reference sequence to identify their positions, motifs and flanking sequences. Twelve STRs from our STR reference panel had matching coordinates and motif sequences. Among them, 10 STRs were genotyped in 183 samples with allele counts ranging from a minimum of 5 to a maximum of 11, whereas the remaining 2 STRs were excluded during quality filtering.

Functional annotation showed an enrichment of STRs in intergenic regions (60.4%, p=0.002, OR=1.26). STRs were depleted in exonic regions (0.89%, p=0.003, OR=0.36) and promoter regions (3.55%, p=0.027, OR=0.66) (Figure 1d & Figure S7 in File S1). The proportion of STRs that overlapped exons was highest for tri- (1.9%) and hexanucleotide (1.4%) motifs (Figure 1d) which were the least heterozygous among all annotation categories (Figure S8 in File S1).

### Structural variant discovery and genotyping

We applied the smoove workflow to discover and genotype 61,806 SVs in the 183 cattle genomes, of which we retained 19,408 polymorphic autosomal loci (12,899 deletions (DEL), 1,043 duplications (DUP), 224 inversions (INV) and 5,242 SVs with unspecified breakends (BND)) after stringent filtering for downstream analyses (Figure 2a in File S1 and Table S5 in File S2). The number of polymorphic SVs identified per chromosome was correlated (r=0.94) with chromosome length (Figure S9 in File S1). We found between 4,259 and 6,835 SVs in each cattle genome (mean: 5,915), and this number was correlated with sequencing coverage (r=0.60) (Figure S10 in File S1). A total of 6,728 (34.6%) SVs had minor allele frequency below 0.05 (Figure 2c). Inspecting the length of the different SV types suggested that most (n=891, 85.4%) DUP were smaller than 1 kb, whereas 3,465 (23%) DEL were larger than 1 kb (Figure S11 in File S1).

**Figure 2:**
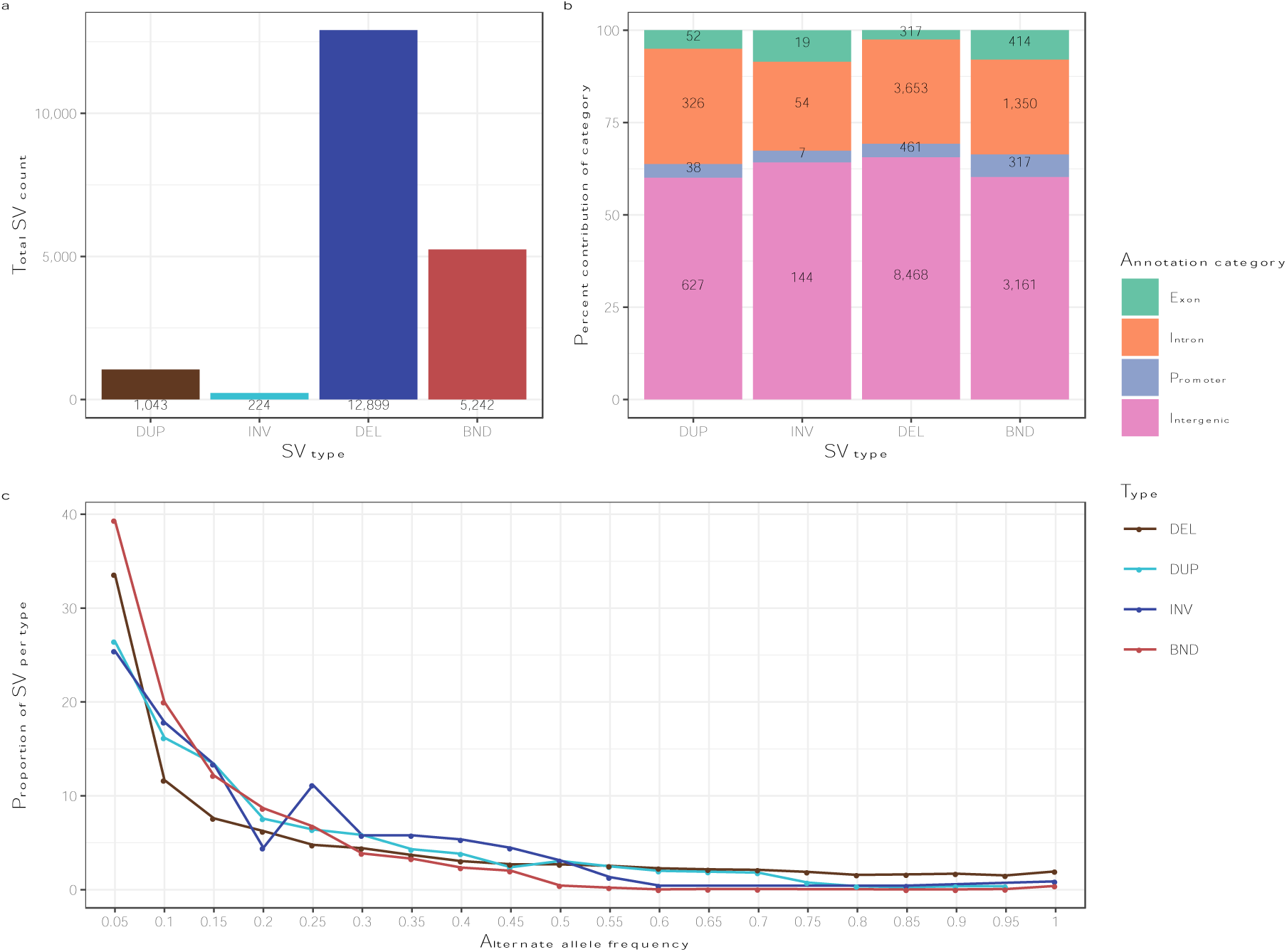
Properties of 19,420 polymorphic SVs in 183 taurine cattle genomes. (a) Count of polymorphic loci for each SV type. (b) Proportion of loci overlapping four annotation categories. The numbers inside the stacked bars represent the count of SVs for each annotation category. (c) Alternative allele frequency distribution for each SV type.

We annotated the SVs according to their location to assess putative functional consequences. This approach revealed that 12,400 (63.8%), 5,383 (27.7%), 823 (4.2%) and 802 (4.1%) SVs overlapped intergenic, intronic, promoter and exonic regions, respectively (Figure 2b). SVs partially or fully overlapped 3,863 genes. Among the SVs that overlapped exons, we identified 52 DUP (25 copy gain DUP or whole gene DUP, 4 full exonic DUP and 23 partial exonic DUP), 19 INV (15 whole gene INV and 4 INV with one breakpoint in exon) and 317 putative loss of function DEL (162 whole gene DEL and 155 DEL affected at least one exon with one breakpoint). We also detected 414 BND in exons. Whole gene inversions (median size 867.9 kb) were the type of exonic SV that was largest in size and lowest in number. The whole-gene inversions detected encompassed 182 coding genes and 32 non-coding genes. Approximately one third of the SVs (n=7,083, 36.4%) were only present in the heterozygous state, and most of these (n=4,989, 70.4%) had minor allele frequency less than 0.05 (Figure 2c). Among these, 4,345, 2,017, 391 and 330 overlapped with intergenic, intronic, promoter, and exonic regions, respectively.

### Linkage disequilibrium and population structure of the cattle cohort

We also discovered and genotyped SNPs and Indels in the 183 animals using the GATK haplotype caller. We considered 12,222,397 SNPs and 1,317,363 Indels with minor allele frequency greater than 5% for the downstream analyses, of which 55,010 SNPs and 89,673 Indels overlapped with STRs, and 387,593 SNPs and 47,129 Indels overlapped with SVs. The large overlap between SNP, SV and STR variants is possibly due to nested variation but can also indicate that short sequencing reads are unable to resolve complex DNA variation.

We calculated the principal components from genomic relationship matrices built with SNP, STR and SV genotypes of the 183 cattle. All three analyses correctly separated the individuals by breed (Figure S12 in File S1). Due to variation in sample size, coverage, and insert size between breeds, we did not investigate within- and across-breed diversity in SVs and STRs. Next, we investigated if SVs and STRs can be tagged by SNPs/Indels. We calculated the linkage disequilibrium (LD) between SNPs/Indels within 100 kb of each SV and STR. We observed that 40.1% of STRs (n=150,393) were in high LD (R^2^ > 0.8) with at least one SNP or Indel while this fraction ranged from 3.1% and 52.2 % for the different SV types (Figure S13a in File S1 and Table S6 in File S2). BND and DUP were poorly tagged, possibly indicating low genotyping accuracy for these loci. The LD between SNPs/Indels and STRs was consistent across the different STR types (Figure S13b and Supplementary Table S6 in File S2).

### Properties of STRs and SVs associated with gene expression

The impact of polymorphic SVs, STRs, SNPs and Indels on gene expression was investigated in a subset of 75 sequenced bulls that also had testis RNA sequencing data. We performed cis-eQTL mapping between 19,415 expressed genes and 12,093 SVs, 271,450 STRs and 13,494,075 SNPs and Indels that had minor allele frequency greater than 5% in the 75 bulls. Five eQTL analyses were conducted, i.e., for SNPs & Indels, SVs, STRs, and jointly for SVs and STRs (SV-STR), and all (ALL) variants to assess the contribution of different types of DNA variation to gene expression.

An eQTL mapping with 13,494,075 ALL variants revealed 6,627 eGenes associated with 7,398 unique eVariants (25 SVs, 514 STRs, 964 Indels & 5,902 SNPs). Both SVs (OR=3.98 and p=1.4 × 10^−8^) and STRs (OR=3.6, p=6.7 × 10^−125^) were enriched among the eVariants indicating that these variant types contribute disproportionally to gene expression variation. The SV-STR eQTL mapping revealed 5,641 eGenes associated with 5,971 unique eVariants (Table 1). The subsequent separate variant type eQTL mapping revealed 6,550, 1,798 and 5,669 eGenes with 7,303, 1,391, 5,995 unique eVariants, respectively, when only SNPs/Indels, SVs and STRs were considered (Table 1). A total of 1,514 eGenes overlapped between the five eQTL analyses (Figure S14 in File S1). Most eGenes (3,379) were shared between the separate eQTL analyses but 1,420 eGenes were shared only between SNPs & Indels and ALL suggesting that many eGenes are only associated with SNPs and Indels. A larger proportion of eSV (24.1% of eSV) and eSTR (8.0% of eSTR) than eSNV/eIndel (2.9% of eSNP/Indel) were associated with the expression of multiple eGenes (Table 1).

**Table 1:** Overview of cis-eQTL detected in 75 testis transcriptomes.

| Type | Total variants | eGenes | e-Variant (% total variants) | eVariant (affecting > 1 eGene) | eQTL (eVariant-eGene pair) |
| --- | --- | --- | --- | --- | --- |
| SNP & Indel | 13,210,530 | 6,550 | 7,303 (0.05%) | 153 | 7,665 |
| SV | 12,093 | 1,798 | 1,391 (11.5%) | 336 | 1909 |
| STR | 271,450 | 5,669 | 5,995 (2.2%) | 485 | 6,572 |
| SV-STR | 283,545 | 5,641 | 5,971 (2.1%) | 465 | 6,525 |
| ALL | 13,494,075 | 6,627 | 7,398 (0.05%) | 146 | 7,552 |

The contribution of STRs and SVs to gene expression variation was quantified based on results from the SV-STR eQTL analysis (Table S7 in File S2). The eVariants were more strongly enriched for SVs than STRs (312 eSV, OR=1.2, p = 3.3 × 10^−4^) but most eVariants were STRs (5,659 eSTRs out of 5,971 eVariants). Among the different SV types, DEL were enriched (264 eDEL, OR=1.6, p=2.9 × 10^−12^) and BND were depleted (30 eBND, OR=0.4, p=3.3 × 10^−7^) among the eVariants compared to STRs (Figure 3a and Table S8 in File S2). The proportion of eVariants associated with multiple eGenes was higher for eDUP (21.4%) than eSTRs (9.6%). Overall, eDUP affected on average 1.35 eGenes (eSV 1.12 eGenes) whereas eSTR and eSNP & eIndel affected 1.11 and 1.01 eGenes, respectively. The maximum number of eGenes per eVariant was larger for STR (n=6) than any other variant type (Figure 3b).

**Figure 3:**
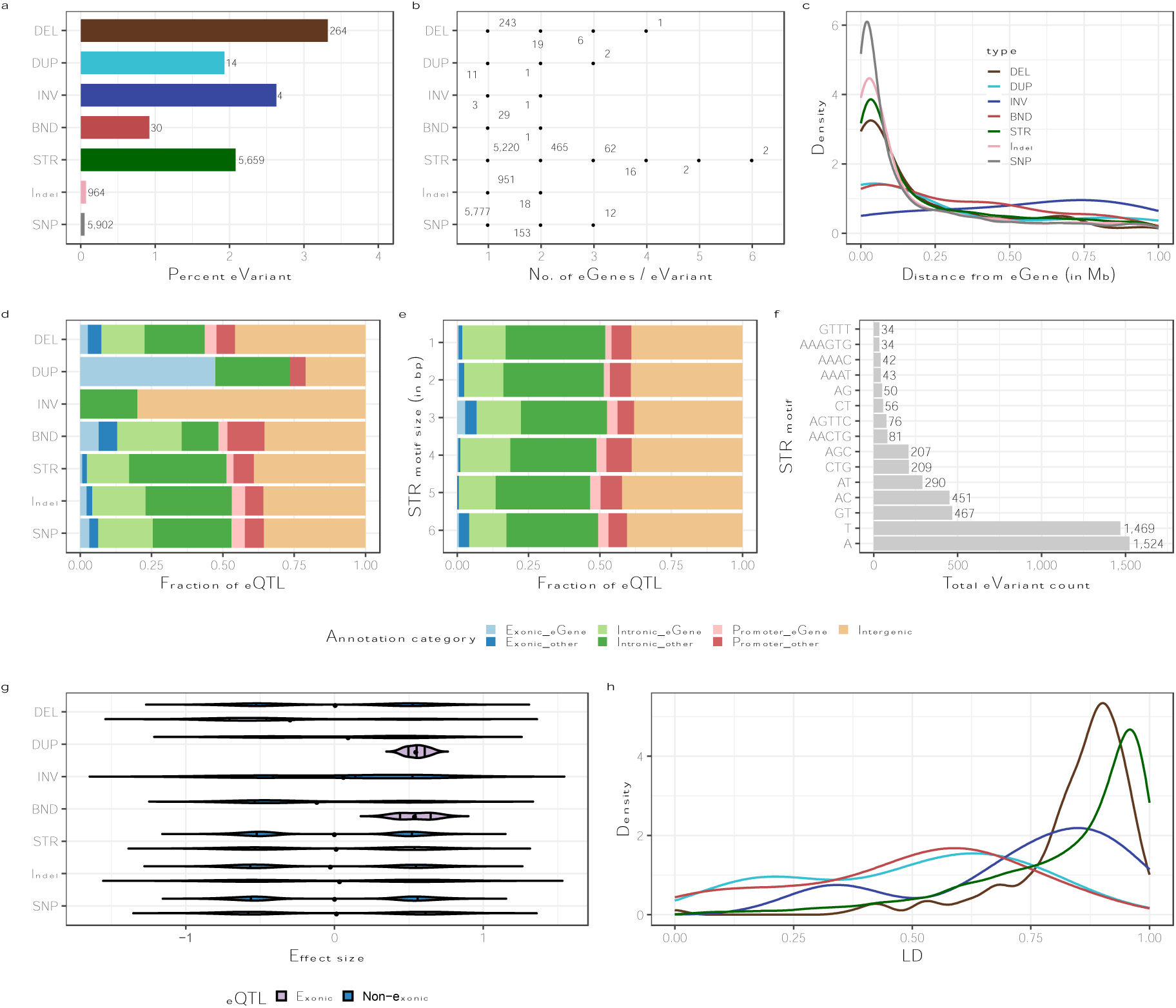
Properties of eSVs and eSTRs from the SV-STR eQTL mapping. eSNPs and eIndels are added from the ALL eQTL mapping. (a) Percentage of unique eVariants for each variant type. The count of eVariants per type is shown next to each bar. (b) Number of eGenes affected by each type of eVariants. (c) Distribution of the absolute distance between eVariants and eGenes (5’UTR or TSS). (d) Proportion of eQTLs from different annotation categories in each variant type, such as those overlapping intergenic regions or overlapping exons, promoters, or introns of their corresponding eGenes or other genes. (e) Proportion of eQTLs from different annotation categories in each STR type such as those overlapping intergenic regions or overlapping exons, promoters, or introns of their corresponding eGenes or other genes. (f) Total count of the most frequent STR motifs (>30 observations) among eSTR. (g) Distribution of effect size of eQTL per type based on exonic (overlap with an exon (exonic) of their eGene or other genes) or non-exonic category. (h) Distribution of maximum LD (R^2^) per variant for each eVariant type.

We examined the distance between eVariants and eGenes (5′-UTR or TSS) and found that most eSVs and eSTRs were located within 250 kb of eGenes (Figure 3c) but eBND (48.3%) and eINV (80%) were more distant (>250 kb) from their eGenes (Table S9 in File S2). Overall, 19.9% eVariants (n=1,194) overlapped with their eGenes; 64 (0.9%), 166 (2.5%) and 964 (14.7%) overlapped with exons, promoters, and introns. Most eQTL were in introns (48.1%) or intergenic regions (39.4%). eDUP were enriched in exons of their eGenes (OR=105.1, p=4.0 × 10^−4^) while eDEL were enriched in the exons of other genes (OR=2.89, p=9.2 × 10^−4^). In contrast, eSTRs were depleted in the exons of their eGenes (OR=0.1, p= 3.4 × 10^−4^) and other genes (OR=0.4, p=6.2 × 10^−4^) (Figure 3d and Table S10 in File S2). These results suggest that eDUP impact gene expression by increasing the copy of their eGenes which agrees with previous research (Shaul 2017). The highest proportion of eSTRs overlapping with exons of their eGenes (17 eSTR) or other genes (24 eSTR) had a trinucleotide repeat motif (Figure 3f). Such STRs are likely to be more tolerated and less selected against than those compromising the triplet codon structure. Most trinucleotide eSTRs in exons had GC-rich repeat motifs (CGG, CTG, CCG, AGC).

Exonic eDUP predominantly increased gene expression, while exonic eDEL mostly decreased gene expression. All other eVariant types exhibited a bimodal effect size distribution. We then explored the LD between eSTRs and eSVs and nearby SNP/Indel. More than three quarter (78.4%) of the eDEL and two thirds of the eSTR (65.9%) were in high LD (R^2^> 0.8) with surrounding SNP/Indel. In contrast, eDUP and eBND were poorly tagged by SNP/Indel.

We found that 92.2% of the eGenes were protein-coding genes, 4.5% were lncRNA, 0.98% were pseudogenes, and 1.8% were other genes (i.e., genes which are not classified in the above three categories) (Figure S15 in File S1). We observed a similar distribution of eGenes across all eVariant types except for eINV.

We identified a candidate causal eSTR (GT_11_, Bos_Tau_STR_126581, Chr10:102,255,360– 102,255,381 bp) in the seventh intron of *TTC7B* encoding tetratricopeptide repeat domain 7B (Figure 4a). The abundance of *TTC7B* mRNA (mean TPM 4.9 ± 1.3) increased with an expansion of the GT repeat motif (p=3.4 × 10^−12^). This STR was the top eVariant in both the ALL and SV-STR eQTL analyses (Figure 4b, c). A candidate causal eSV is a 885 bp deletion (Chr4:99,481,913–99,482,798 bp) encompassing *ENSBTAG00000015551* and the distal end of *SLC13A41* encoding solute carrier family 13-member 4 (Figure 4d). The deletion reduced mRNA expression of *ENSBTAG00000015551* (mean TPM 7.2 ± 3.4, p=7.1 × 10^−17^) and *SLC13A4* (mean TPM 0.9 ± 0.4, p=1.6 × 10^−15^) (Figure e,f). This deletion was the top eVariant in the ALL and SV-STR eQTL analyses for both genes.

**Figure 4:**
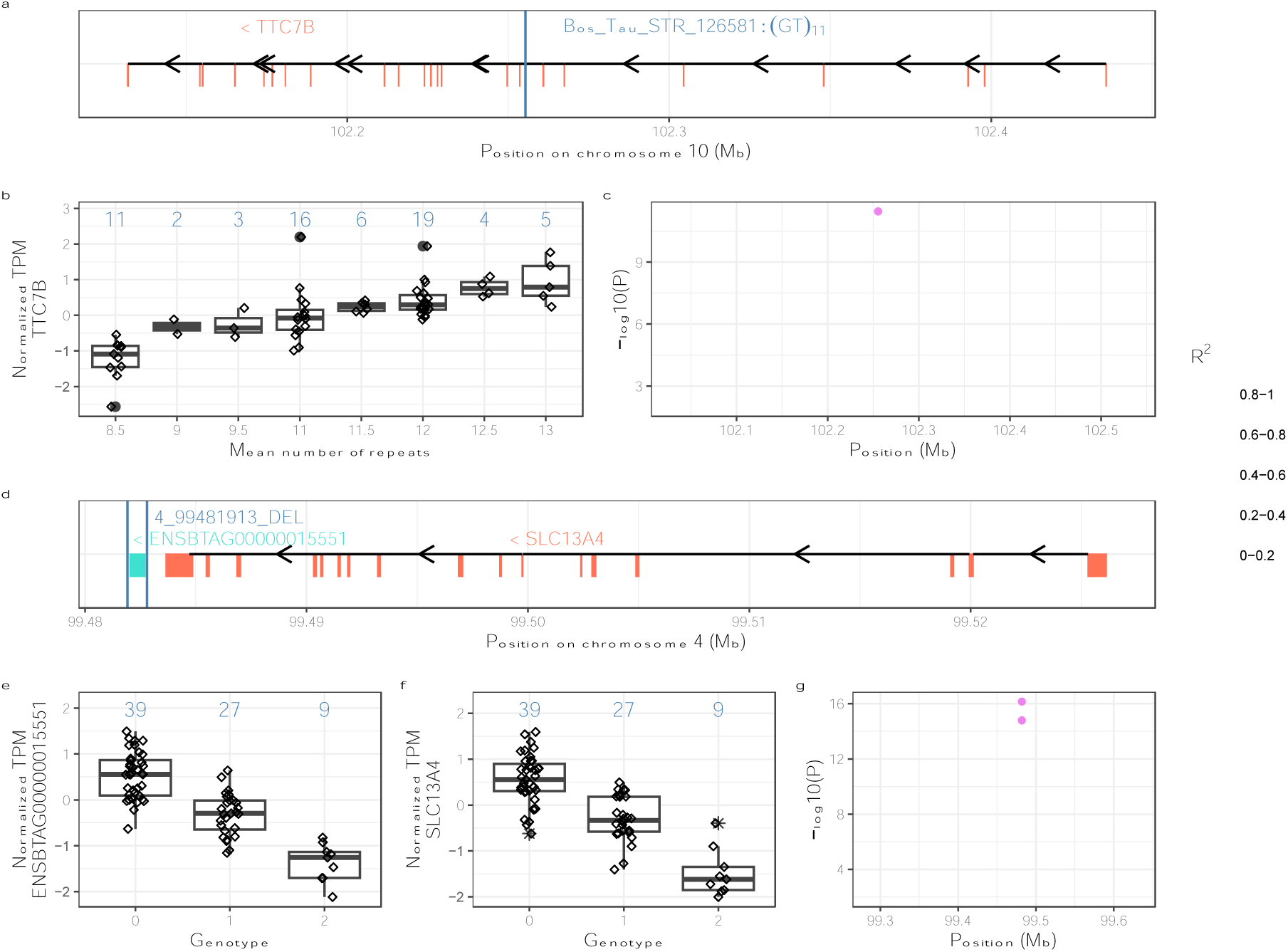
Candidate eSTR (a-c) and eSV (d-g) associated with eGene expression. The eSTR Bos_Tau_STR_126581 is a GT dinucleotide that repeats 11 times in the reference sequence, and between 8.5 and 13 times in the 75 genotyped bulls. eQTL mapping revealed association between Bos_Tau_STR_126581 and *TTC7B* mRNA abundance. (a) Schematic overview of the exon/intron structure of bovine *TTC7B* gene and Bos_Tau_STR_126581. The vertical blue line indicates the position of Bos_Tau_STR_126581 and the vertical salmon lines indicate exons of *TTC7B*. (b) Normalized gene expression of *TTC7B* in 75 genotyped bulls in each mean dosage of eSTR. The mean repeat count per genotype is determined by adding the total repeat number of both alleles belonging to a genotype and then dividing this sum by 2. (c) Manhattan plot of –log10(P)-values for all variants surrounding Bos_Tau_STR_126581 from the nominal ALL-eQTL analysis. Different colours indicate the pairwise linkage disequilibrium (R^2^) between Bos_Tau_STR_126581 and all other variants. (d) Schematic overview of *ENSBTAG00000015551* (turquoise colour) and *SLC13A41* (salmon colour) that are associated with a 885 bp deletion on chromosome 4 (eDEL 4_99481913_DEL). The boxes represent exons. The vertical blue lines indicate the position of 4_99481913_DEL. (e) & (f) Normalized mRNA expression of *ENSBTAG00000015551* and *SLC13A41* in 75 genotyped bulls for each genotype of eDEL. The blue numbers are the total number of animals per genotype. (g) Manhattan plot of –log10(P)-values for all variants surrounding 4_99481913_DEL from the nominal ALL-eQTL analysis as pink colour (two points as same eDEL corresponds to two genes). Different colours indicate the pairwise linkage disequilibrium (R^2^) between 4_99481913_DEL and all other variants.

### Cis-sQTL mapping

We calculated intron excision ratios of 241,427 introns assigned to 76,083 intron clusters. More than half (n=135,342, 56.0%) of the introns overlapped with 14,583 genes, but the annotation-free splicing event identification by the LeafCutter software also detected many introns that did not overlap with annotated features. The intron excision ratios were normalized for each intron and subsequently used as input phenotypes for cis-sQTL mapping. We mapped cis-sQTL with an approach that was similar to the eQTL mapping, i.e., we separately considered SNPs & Indels, SVs, STRs, SV-STR, and ALL.

The ALL sQTL mapping revealed association between 12,835 unique lead variants (sVariant) and 11,588 (15.2%) intron clusters (sIntron cluster). The 12,835 sVariants included 25 SVs, 712 STRs, 1,593 Indels & 10,505 SNPs, and 286 of the sVariants were associated with more than one intron cluster. More than half of the sIntron clusters (n=6,798, 58.6%) overlapped with 4,890 sGenes whereas the remaining did not overlap with annotated features. Both SVs (OR=2.3, p=2.3 × 10^−4^) and STRs (OR=2.9, p=6.4 × 10^−123^) were enriched among the sVariants when compared to SNPs and Indels. The SV-STR analysis revealed 9,065 sIntron clusters associated with 8,857 unique sVariants (Table 2). Variant type-specific sQTL analyses revealed 8,749, 1,707 and 12,683 sVariants, respectively, when only STR, SV and SNP & Indel were considered (Table 2).

**Table 2:**
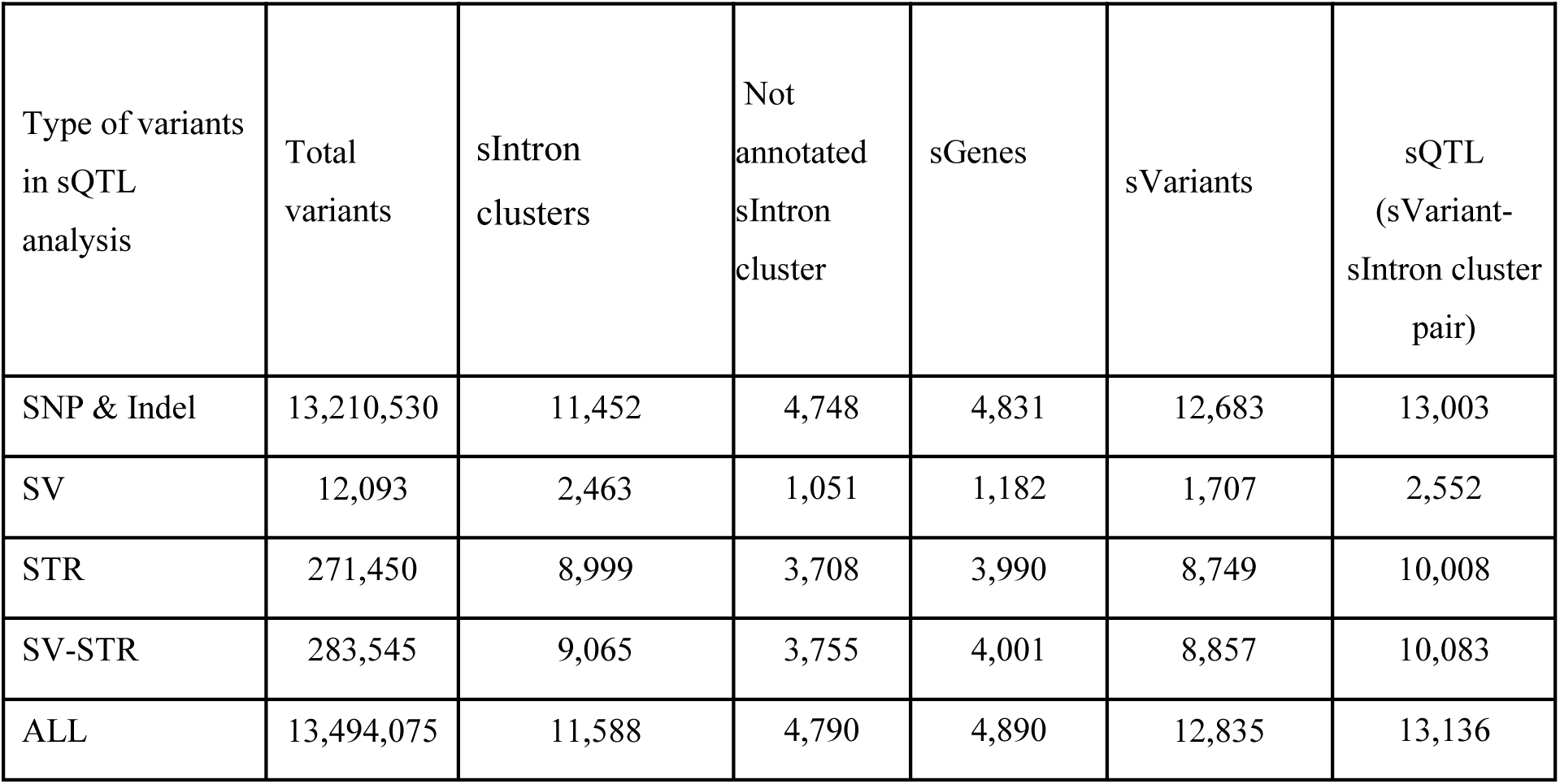
Overview of cis-sQTL detected in 75 testis transcriptomes.

| Type of variants in sQTL analysis | Total variants | sIntron clusters | Not annotated sIntron cluster | sGenes | sVariants | sQTL (sVariant-sIntron cluster pair) |
| --- | --- | --- | --- | --- | --- | --- |
| SNP & Indel | 13,210,530 | 11,452 | 4,748 | 4,831 | 12,683 | 13,003 |
| SV | 12,093 | 2,463 | 1,051 | 1,182 | 1,707 | 2,552 |
| STR | 271,450 | 8,999 | 3,708 | 3,990 | 8,749 | 10,008 |
| SV-STR | 283,545 | 9,065 | 3,755 | 4,001 | 8,857 | 10,083 |
| ALL | 13,494,075 | 11,588 | 4,790 | 4,890 | 12,835 | 13,136 |

We then assessed the overlap of sGenes/sIntron clusters between all sQTL analyses. Approximately half of the sIntron-clusters (n=6,034, 47.9%) and sGenes (n=2,590, 49.1%) overlapped between the SNPs & Indels, STR, SV-STR and ALL sQTL analyses suggesting that distinct variant types in LD tag the same splicing event (Figure S16 and S17 in File S1). A total of 3,126 (24.8%) sIntron clusters and 1,126 (21.3%) sGenes were shared only between SNPs & Indels and ALL suggesting that a substantial fraction of sGenes is only associated with SNPs and Indels.

### Variant properties of sSTR and sSV

The impact of STRs and SVs on alternative splicing was assessed based on the results from the SV-STR sQTL analysis (Table S11 in File S2). We observed that DEL were more likely to be sVariants than STRs (4.9% of DEL, OR=1.6, p=1.7 × 10^−17^) (Figure 5a and Table S12 in File S2). Conversely, BND (OR=0.4, p=1.3 × 10^−9^) were less likely to be sVariants compared to STRs. We further examined how many intron clusters are affected by an sVariant. A similar proportion of sDEL (12.2%) and sSTRs (11.2%) were associated with multiple sIntron clusters whereas this fraction was considerably lower or negligible for all other types of sVariants (Figure 5b). Between 62% and 81% of the sVariants were located within 100 kb of their sIntron cluster (Figure 5c and Table S13 in File S2).

**Figure 5:**
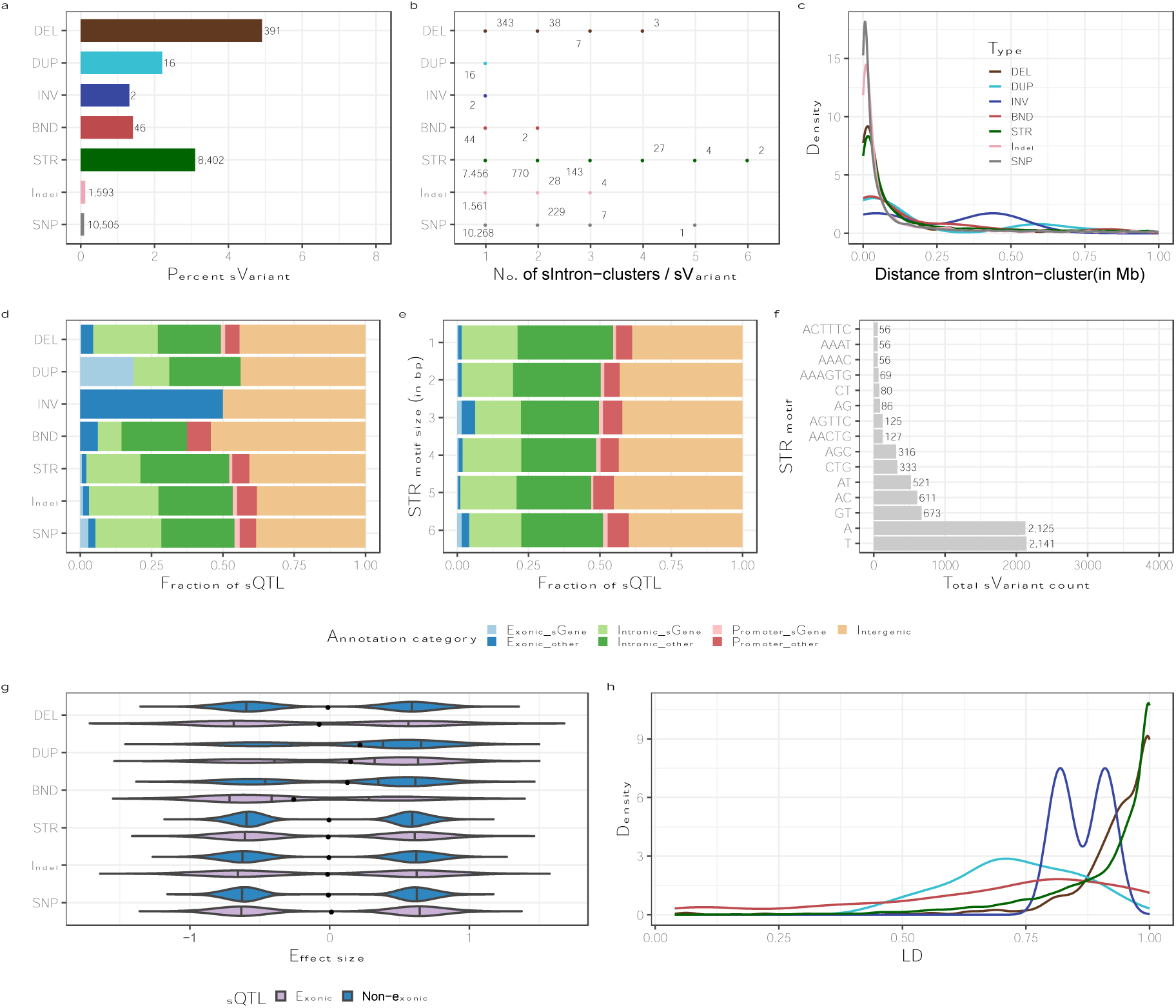
Properties of sVariants (SVs and STRs) from the SV-STR sQTL analysis. SNPs & Indels are from ALL analyses in all panels (a) Percentage of unique sVariants for each variant type. The number of sVariants per category is shown next to the bars. (b) Total number of sIntron clusters per sVariant for each variant type. (c) Distance between sQTL and the start position of the associated sIntron cluster for each variant type. (d) Fraction of sQTL from different annotation categories in each variant type such as those overlapping intergenic regions or overlapping exons, promoters, or introns of their corresponding sGenes or other genes. (e) Fraction of sQTL from different annotation categories in each STR motif size such as those overlapping intergenic regions or overlapping exons, promoters, or introns of their corresponding eGenes or other genes. (f) Prevalence of the most frequent (>50) STR motif among sSTR. (g) Distribution of sQTL effects. Colours differentiate between exonic and non-exonic sQTL. The black dots represent the overall mean. (h) Distribution of maximum LD (R^2^) between sQTL and SNP/Indel for different variant types.

Most of the sQTL overlapped with either introns (49.7%) or intergenic regions (41.0%), but only few with promoter (6.9%) and exons (2.3%). Interestingly, sDEL were enriched in exons and depleted in introns of other genes, whereas sDUP showed enrichment in exons of sGenes (Table S14 in File S2). On the other hand, sSTRs were depleted in exons of other genes but they were enriched in introns of other genes (Table S14 in File S2). We observed a high proportion of trinucleotide sSTRs among those that overlapped exons. These trinucleotide sSTRs were GC-rich (Figure 5f). Most sQTL showed bimodal effects. A bimodal effect size distribution in splicing variation encompassing both positive and negative effects is associated with variation in the relative abundance of transcripts between different genotypes (Garrido-Martín et al. 2021). sDUP had slightly positive effects on splicing phenotypes which may indicate a relatively higher abundance of the transcript associated with the duplication (Figure 5g). The vast majority of sDEL (94.0%) and sSTRs (84.3%) were in high LD (R^2^> 0.8) with surrounding (±50 Kb) SNP/Indel, but sDUP (31.5%) and sBND (39.5%) were less frequently tagged (Figure 5h).

Finally, we compared genes and molecular QTL (eQTL and sQTL as gene-variant pair) from both the eQTL and sQTL SV-STR analyses (Figure S18 in FileS1). This comparison revealed that 1,988 genes and 505 QTL overlapped between both analyses. Out of the 505 shared QTLs, 479 were due to STR, while 23 were due to DEL (Figure S19 in FileS1). The eQTL that were also sQTL mainly regulated expression due to alterations in gene transcript level abundance, and these changes were mainly modulated by STR and DEL.

Among the sQTL, we identified a candidate causal sSTR (AACTG_5_, Bos_Tau_STR_57388, Chr1:112307866–112307890 bp) upstream of the long non-coding RNA (lncRNA) *ENSBTAG00000054182* (Figure 6a). An expansion of Bos_Tau_STR_57388 (the inserted motif AAATG differed slightly from the reference motif AACTG) was associated with a splicing junction (Chr1:112,307,410–112,322,799, p=4.5 × 10^−25^) in both SV-STR and ALL sQTL analyses. This splicing junction extends from upstream the lncRNA to the first intron of the lncRNA (Figure 6a) and its intron excision ratio increased with an expansion of the repeat motif (Figure 6c, d). The expression of *ENSBTAG00000054182* (mean TPM 5.1 ± 1.5) decreased with the insertion of an additional repeat unit (Figure 6b). Bos_Tau_STR_57388 was also the top eVariant for *ENSBTAG00000054182* in the SV-STR eQTL analysis (p=2.3 × 10^−15^) but not in ALL eQTL analysis where a SNP (Chr1:112,315,134 bp) in LD (R^2^=0.93) was the top eVariant.

**Figure 6:**
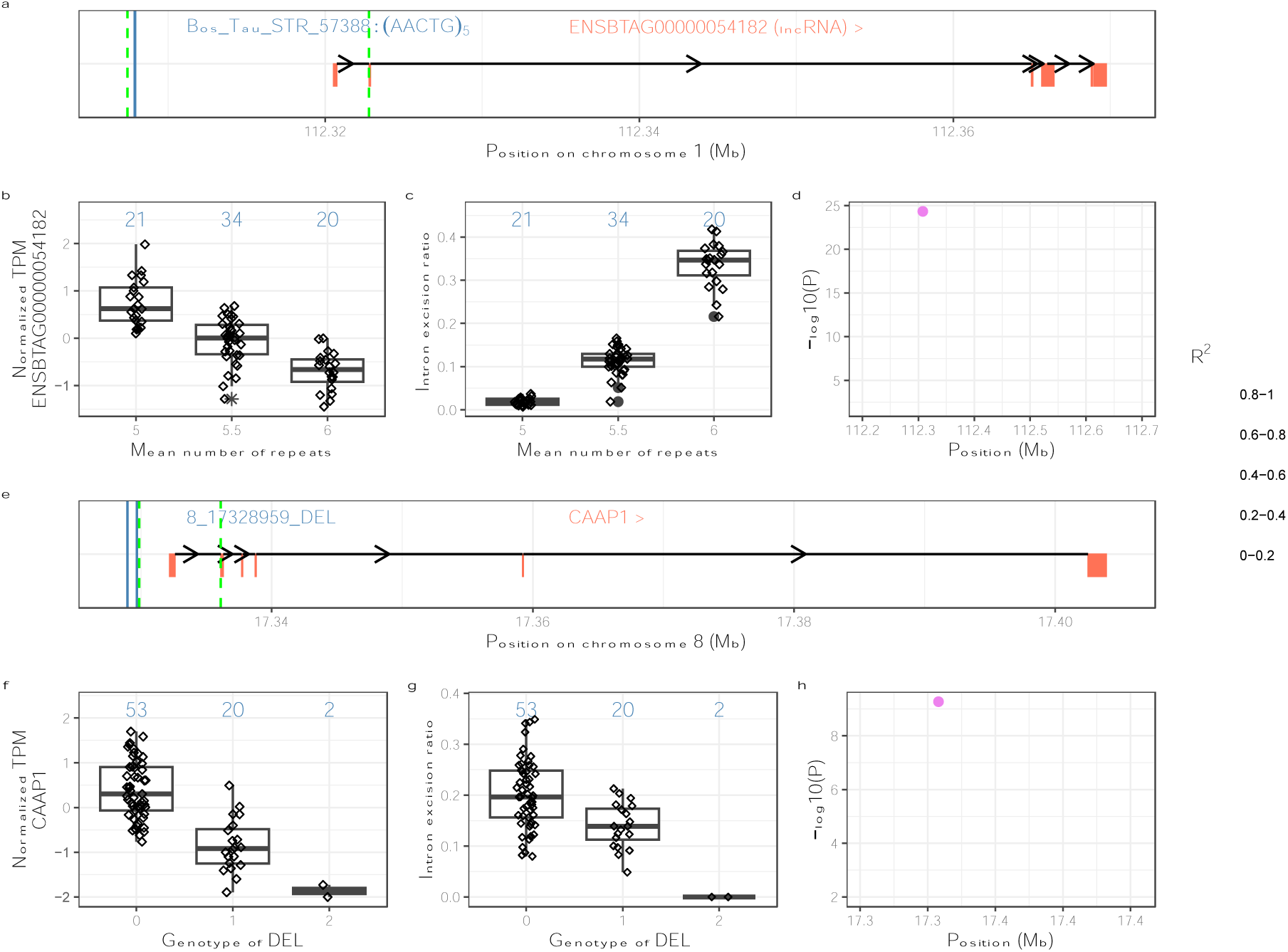
Two candidate causal sSTR (a-d) and sSV (f-h). The sSTR Bos_Tau_STR_57388 on chromosome 1 is associated with *ENSBTAG00000054182* splicing. (a) Schematic overview of *ENSBTAG00000054182.* Bos_Tau_STR_57388 (blue line) is upstream of lncRNA *ENSBTAG00000054182* and it is associated with the splicing junction spanning from Chr1:112,307,410 to 112,322,799. Intron/splice junction boundaries are indicated with green dotted lines. (b) Normalized *ENSBTAG00000054182* expression, (c) intron excision ratio for different sSTR genotypes The mean repeat count per genotype is determined by adding the total repeat number of both alleles belonging to a genotype and then dividing this sum by 2. (d) Manhattan plot of nominal ALL-sQTL result surrounding the sSTR. Different colours indicate the pairwise linkage disequilibrium (R^2^) between Bos_Tau_STR_57388 and all other variants. A Candidate sSV (8_17328959_DEL) on chromosome 8 is associated with alternative *CAAP1* splicing. (e) Schematic overview of *CAAP1* gene. A promoter deletion “8_17328959_DEL” (blue line) is associated with excision ratios of splicing junction Chr8:17,329,855–17,336,097. Intron boundaries are indicated with green dotted lines. (f) Normalized *CAAP1* expression and (g) intron excision ratio for the different sSV genotypes. (h) Manhattan plot of nominal ALL-sQTL result surrounding sDEL. Different colours indicate the pairwise linkage disequilibrium (R^2^) between sDEL and all other variants.

A candidate sSV is a 729 bp deletion on chromosome 8 (Chr8:17,328,959–17,329,688), which resides in the promoter region of *CAAP1* encoding caspase activity and apoptosis inhibitor 1 (Figure 6e). The deletion was associated (p=2.8 × 10^−11^) with reduced excision ratios of a splicing junction (Chr8:17,329,855–17,336,097) overlapping *CAAP1* (Figure 6f, h) and expression of *CAAP1* (mean TPM 26.4 ± 2.8) (Figure 6b). This sDEL was also the top eVariant for *CAAP1* in the ALL eQTL and SV-STR eQTL (p=5.1 × 10^−12^).

## Discussion

We generated a catalogue of bovine polymorphic STRs which contain motifs that vary in size, but some may also contain variation between the repeat motifs. A large number of cattle from different breeds enabled us to genotype sixfold more STRs compared to a previous study (374,821 vs. 60,661) that considered only 5 Holstein cattle genomes (Xu et al. 2017). Three quarter of the STRs genotyped in our study were multiallelic, which agrees with previous studies in cattle, pigs and humans (Willems et al. 2014; Xu et al. 2017; Wu et al. 2021). We also genotyped almost 20k SVs. The majority of both SVs and STRs were in introns and intergenic regions likely because coding regions are less tolerant to variants affecting several bases. We also detected SVs and STRs that overlapped exonic regions but half of the exonic SVs were only present in the heterozygous state which may indicate that some of them manifest deleterious phenotypes in the homozygous state. However, even deleterious SVs can persist and increase in frequency over time due to drift or pleiotropic effects and balancing selection, such as a 660 kb deletion in Nordic red cattle [50]. Deleterious SVs in less conserved genes may be evolutionarily less constrained (Mesbah-Uddin et al. 2018). We also observed a high proportion of tri- and hexanucleotide STR in exonic regions possibly suggesting that non-triplet STRs are less tolerated and might be under negative selection (Willems et al. 2014).

We observed more than twice the number of deletions compared to other SV types likely because they are easier to identify from short-read sequencing data (Mahmoud et al. 2019). Only half of the STRs and DELs are in high linkage disequilibrium with SNPs and Indels (R^2^ > 0.8). The LD between SNPs and other types of SVs such as BND, DUP and INV is even lower which could be due to incorrect genotyping, alignment error, or their occurrence in complex regions such as segmental duplications. Thus, the direct genotyping of these variants is required to enable powerful association studies.

In the cattle breeding industry, genomic prediction is routinely applied to estimate breeding values. This estimation typically relies on dense genotypes obtained from SNP arrays or imputed sequence variants but does not consider an exhaustive set of STRs and SVs. Lee et al. (2023) added 372 SVs to the custom content of a frequently applied microarray to enable genotyping them at the population scale (Lee et al. 2023). Our study confirms that such efforts are warranted, as SVs and STRs are only partially tagged by adjacent SNPs. Particularly intriguing are also SVs and STRs which are associated with gene expression and splicing variation, as they potentially impact phenotypes. Considering such SVs and STRs and those exhibiting limited or no LD with genotyped SNPs carries the potential to improve the accuracy of genomic predictions and offer insights into the molecular underpinnings of QTL (Lee et al. 2021; Leonard et al. 2023).

Our results confirm that sequencing coverage and insert size have profound impacts on the genotyping of SVs and STRs (Willems et al. 2017; Lee et al. 2023). We applied stringent filters to retain only high-confidence SVs. This approach likely removed some true large and complex SVs and STRs from our data. Long sequencing reads and pangenome integration enable to reliably detect large and complex SVs and STRs (Leonard et al. 2022; Talenti et al. 2022). However, long read sequencing is still too costly when applied at population scale. Future studies could utilize a combination of long read sequencing and pangenome integration with short read sequencing data to identify and genotype the full spectrum of genetic variants at the population scale (Chaisson et al. 2019; Ebert et al. 2021).

Our study examines impacts of polymorphic SVs and STRs on gene expression and splicing variation in bovine testis tissue. Further research is needed to extend this analysis to additional tissues as gene regulation is predominantly tissue specific (Gamazon et al. 2018). Regardless, our eQTL and sQTL analyses showed that SVs and STRs have profound impacts on the transcriptional profile in testis. We found that each eSV affects on average 1.11 nearby genes with most of this contribution arising from DUP. However, this value is lower than the 1.82 genes in *cis* per eSV reported recently in humans, where major contributions were from multi-allelic copy number variants (mCNV) and DUP (Scott et al. 2021). In our study, CNV are part of the DUP category. This difference likely indicates that our study had less power to detect s/eQTL because our variant catalogue (61,668 SVs in human vs. 19,408 SVs here) and sample size (643 individuals with 48 tissues vs. 75 individuals with one tissue) were considerably smaller. Our results confirm that e/sDUP in exonic and non exonic regions mostly increase gene expression whereas e/sDEL decrease gene expression (Chiang et al. 2017; Scott et al. 2021). An increased expression associated with an e/sDUP is frequently due to either duplication of the entire gene or exon or its regulatory regions. It is possible that some of the eSVs and eSTRs detected in our study are associated with semen quality and male fertility. Further investigations are needed to explore the integration of molecular QTL cohorts and GWAS cohorts through transcriptome-wide association testing. This approach holds the potential to reveal SVs and STRs associated with specific traits (Mapel et al. 2023).

Our analysis showed that most e/sSTRs and e/sSVs were in intronic regions rather than intergenic regions, which contrasts with their overall distribution along the genome. This pattern agrees with the position of human e/sSTRs and e/sSVs (Chiang et al. 2017; Fotsing et al. 2019; Jakubosky et al. 2020). Our study thus confirms the importance of non-coding SVs and STRs in regulating gene expression and splicing (Jakubosky et al. 2020). Intronic and intergenic regions can contain regulatory elements that modulate splicing and gene expression via change in nucleosome positioning, open chromatin structure, RNA-binding protein or DNA methylation (Fotsing et al. 2019; Ho et al. 2020; Vialle et al. 2022). Nearly half of the intron clusters detected in our study could not be annotated with the current cattle annotation (Ensembl 104). The FANTOM5 consortium revealed significant overlap of transcription start sites (TSS) to STR loci which are unassigned to any known genic or enhancer regions in humans (Grapotte et al. 2021). Most of these TSS overlapping with STRs, are responsible for initiating non-coding RNAs in humans. Similarly, a candidate causal sSTR detected in our study was associated with the splicing of the lncRNA *ENSBTAG00000054182*, which produces a transcript that is not included in the current Ensembl annotation. This further emphasises the need for an improved bovine annotation, particularly with respect to non-coding elements of the genome such as lncRNAs. Although the association of expression and splicing variation with STRs and SVs in e/sQTL studies do not necessarily provide the underpinning molecular mechanism of action, these variants contribute significantly to complex trait variation (Xiang et al. 2022).

## Declarations

### Ethics approval and consent to participate

Not applicable.

### Consent for publication

Not applicable.

### Competing interests

The authors declare that they have no competing interests.

## Data Availability

Short paired-end whole-genome sequencing reads from 183 cattle from five breeds and whole genome RNA-sequencing data from 75 cattle are in the ENA database at the study accessions PRJEB28191, PRJEB46995 and PRJNA238491. The primary output files and raw data info including accession numbers of the raw data, Reference STR, Genotyped STR, and SV, are made available through Zenodo https://doi.org/10.5281/zenodo.8274665. All scripts and workflows are available online: https://github.com/Meenu-Bhati/SV-STR.

## Acknowledgements

We would like to express our gratitude to Dr. Maya Hiltpold for supporting the sampling of testis tissue, Dr. Naveen Kadri for helping in data processing and Dr. Alexander Leonard for support in implementing the Smoove workflow. We also thank the Functional Genomics Center Zurich (Dr. Catharine Aquino) for generating DNA and RNA sequencing data.

## Funding

This study was supported by grants from the Swiss National Science Foundation, an ETH Research Grant, Swissgenetics, and the Swiss Federal Office for Agriculture, Bern. The funding bodies were not involved in the design of the study and collection, analysis, and interpretation of data and in writing the manuscript.

## Author contributions

MB and HP conceived the study. MB aligned RNA-seq data, created and performed the analyses workflows, called STRs and SVs, conducted the e/sQTL mapping, all subsequent analyses and drafted the manuscript. XMM sampled tissue and established the mapping cohort and contributed to the e/sQTL mapping. ALV aligned DNA-seq data and performed variant calling for SNPs and Indels. HP interpreted results and contributed to the writing of the manuscript. All authors read and approved the final version of the manuscript.

## Conflict of Interest

None declared.

## Notes

https://doi.org/10.5281/zenodo.8274665

### Competing Interest Statement

The authors have declared no competing interest.

### Summary of Updates

Manuscript revised in response to feedback from Editor and Reviewers.

