## Supplementary material for "Structural variants and short tandem repeats impact gene expression and splicing in bovine testis tissue": FileS1

**Supplementary Figures**

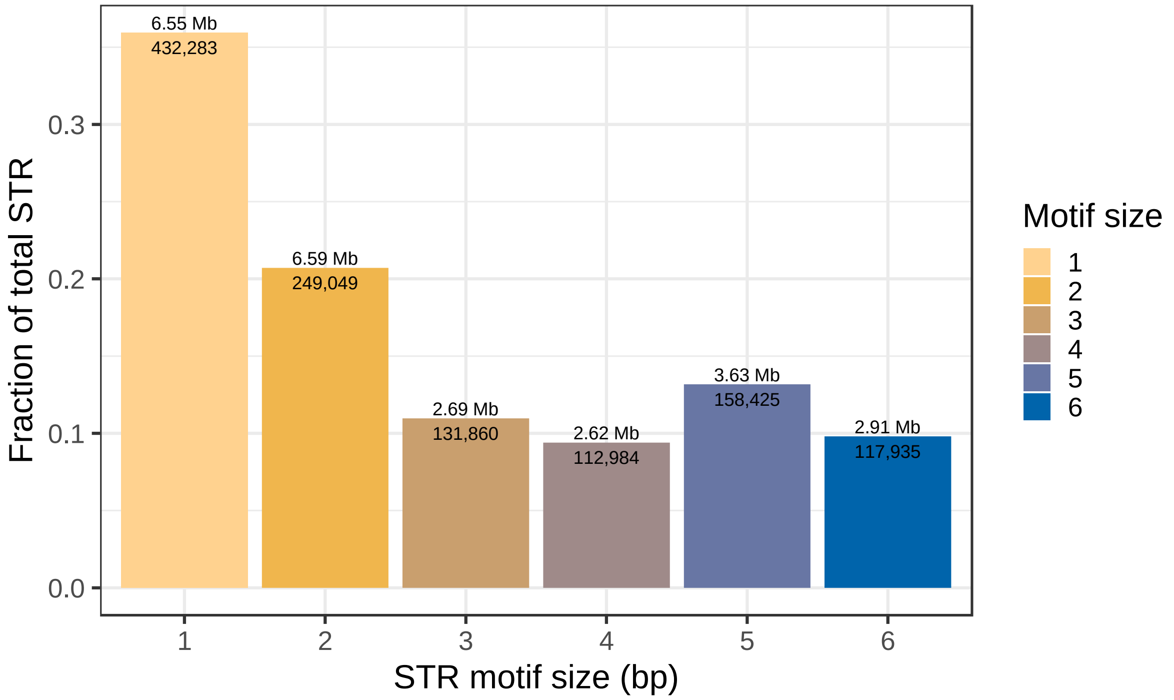


**Figure S1** Proportion and number of STR found in the ARS-UCD1.2 reference sequence in each motif size. The total amount of sequence overlapping each STR motif class is shown above each bar. Total number of loci for each STR motif class is inside the bars. Colours indicate different motif sizes.

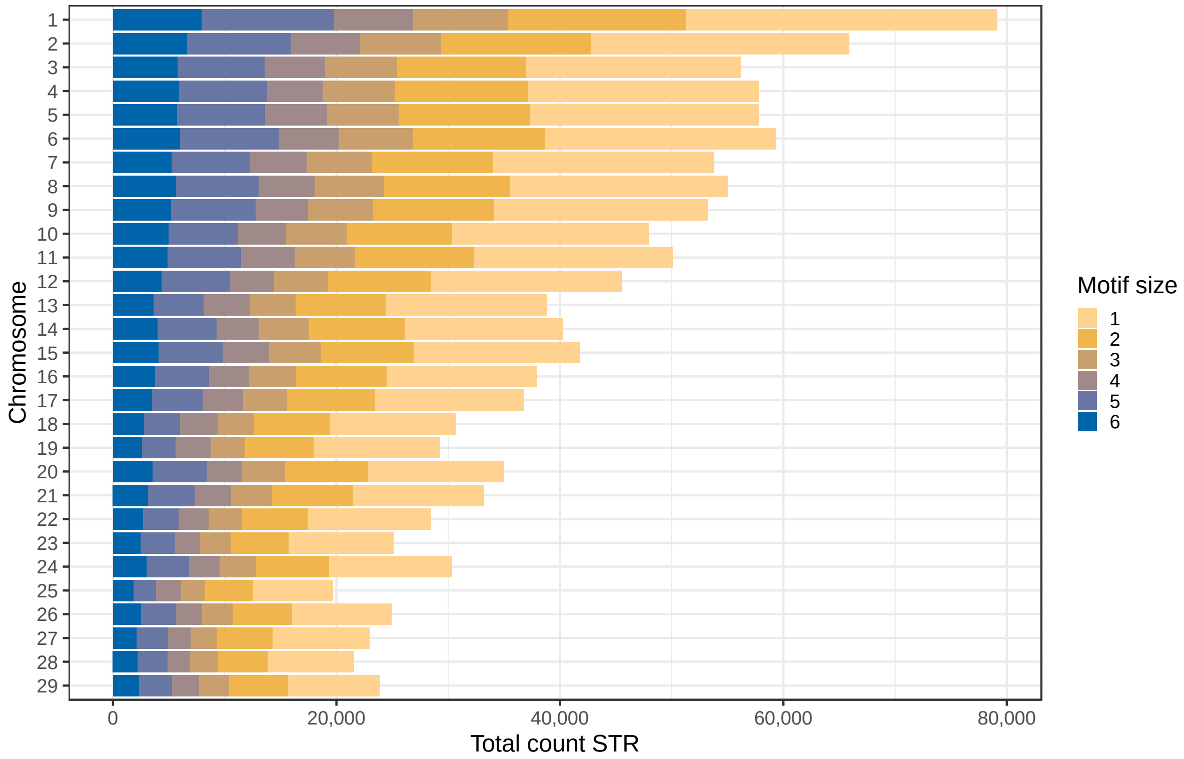


**Figure S2** Cumulative count of reference STR for each autosome. Colours indicate different motif sizes.

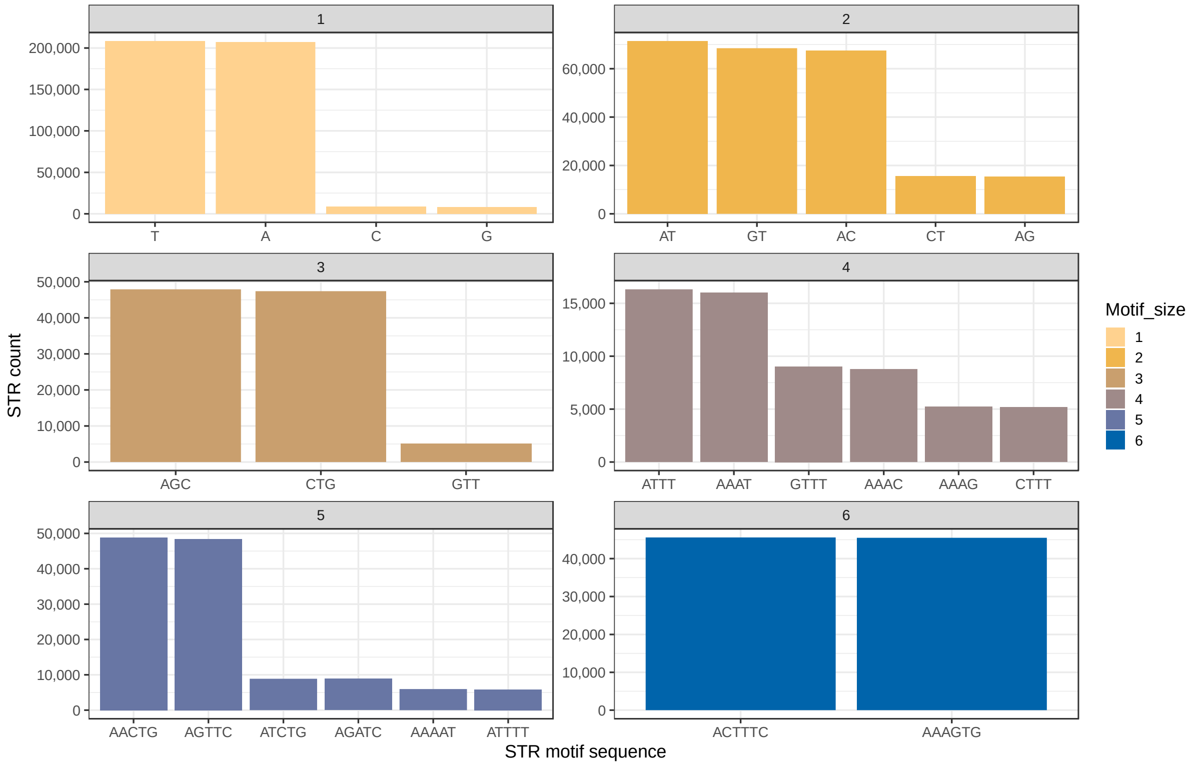


**Figure S3** Most frequent (n>5000 count) motif sequence observed in each STR motif size class in the reference STR. Colours indicate different motif sizes.

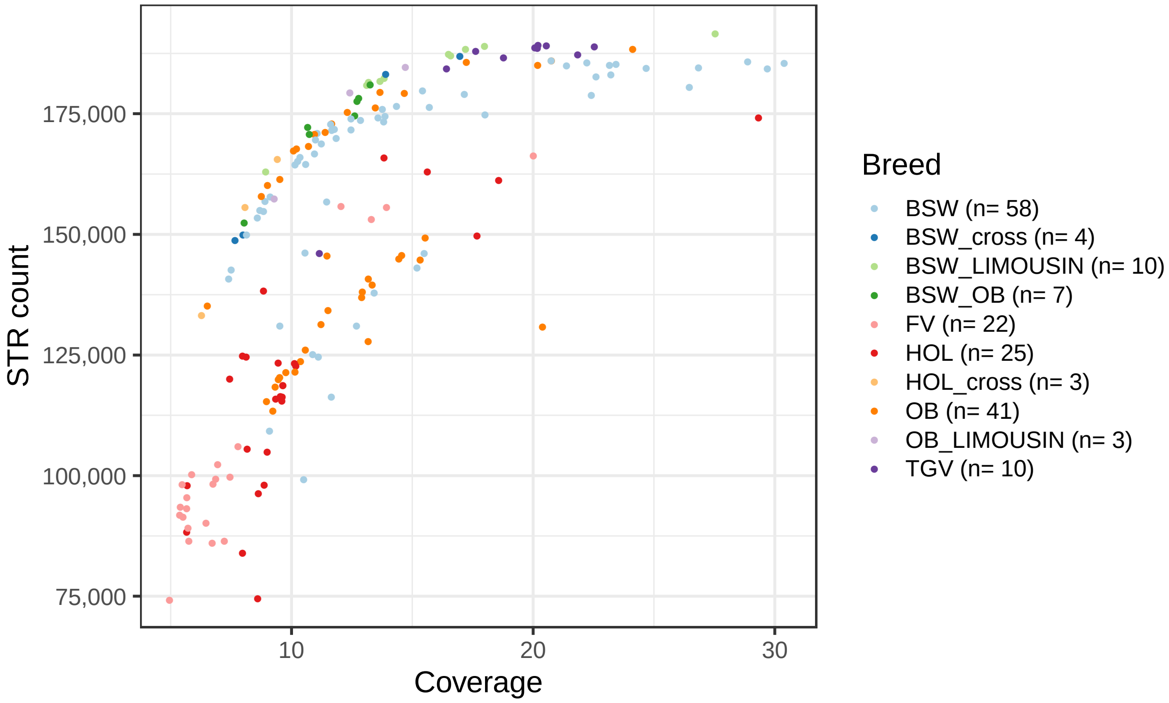


**Figure S4** Number of genotyped STRs and autosomal coverage per animal. Colours represent different breeds.

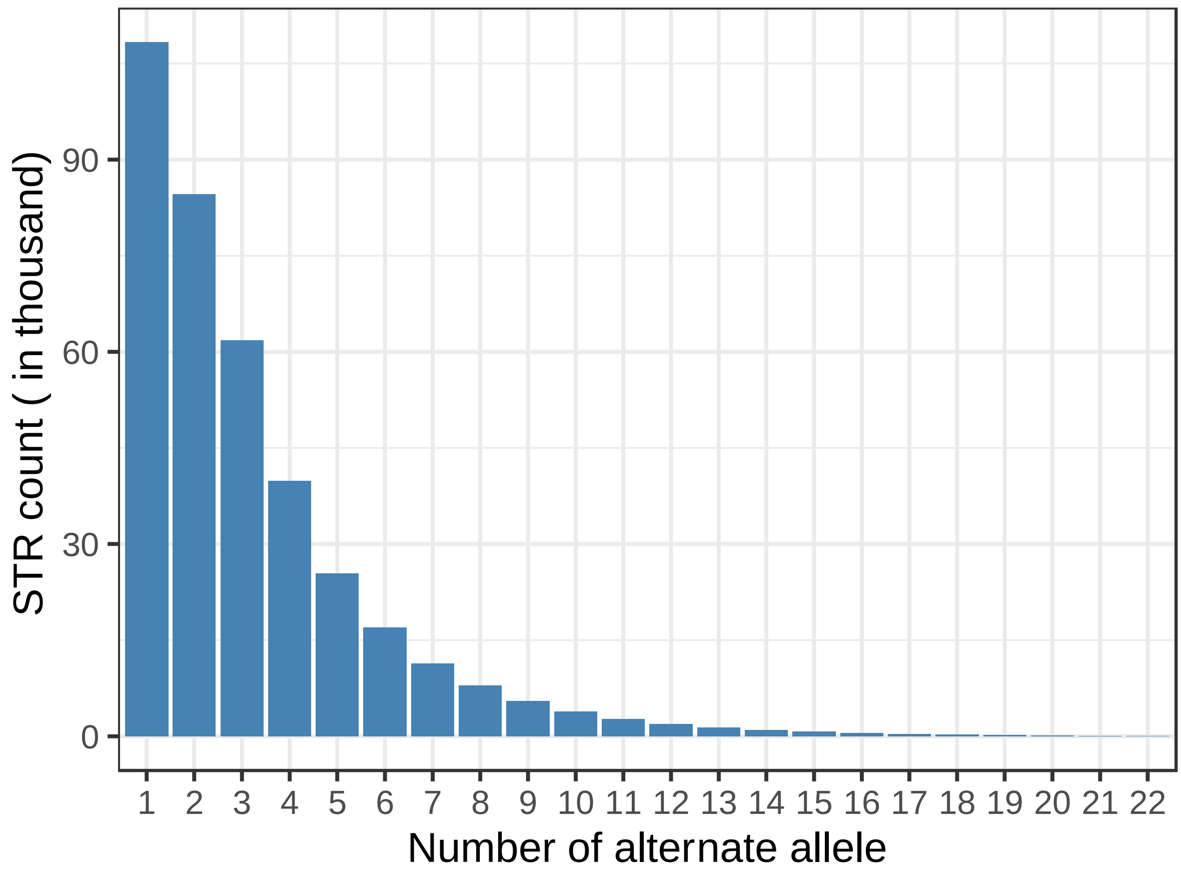


**Figure S5** Number of alternative alleles observed for the genotyped STR. The x axis is truncated at 22.

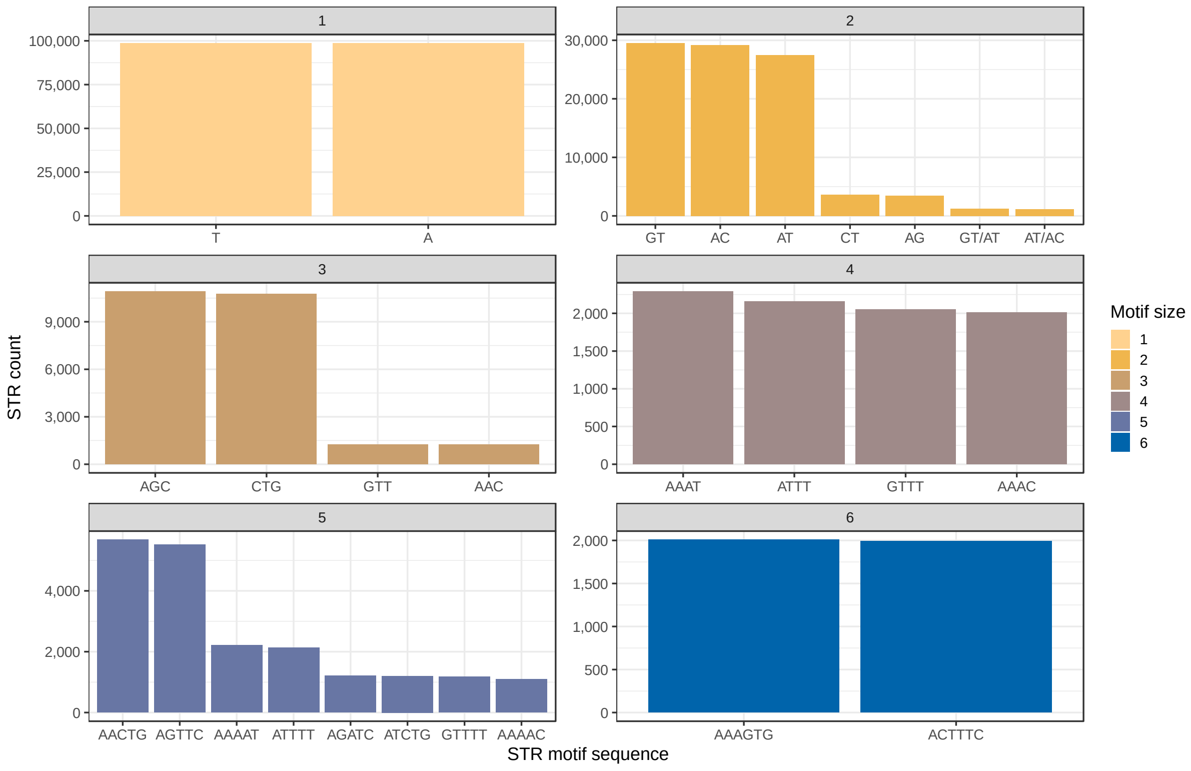

**Figure S6** Most frequent (>1000 count) motif sequence observed in the genotyped STR. Colours indicate different motif sizes.

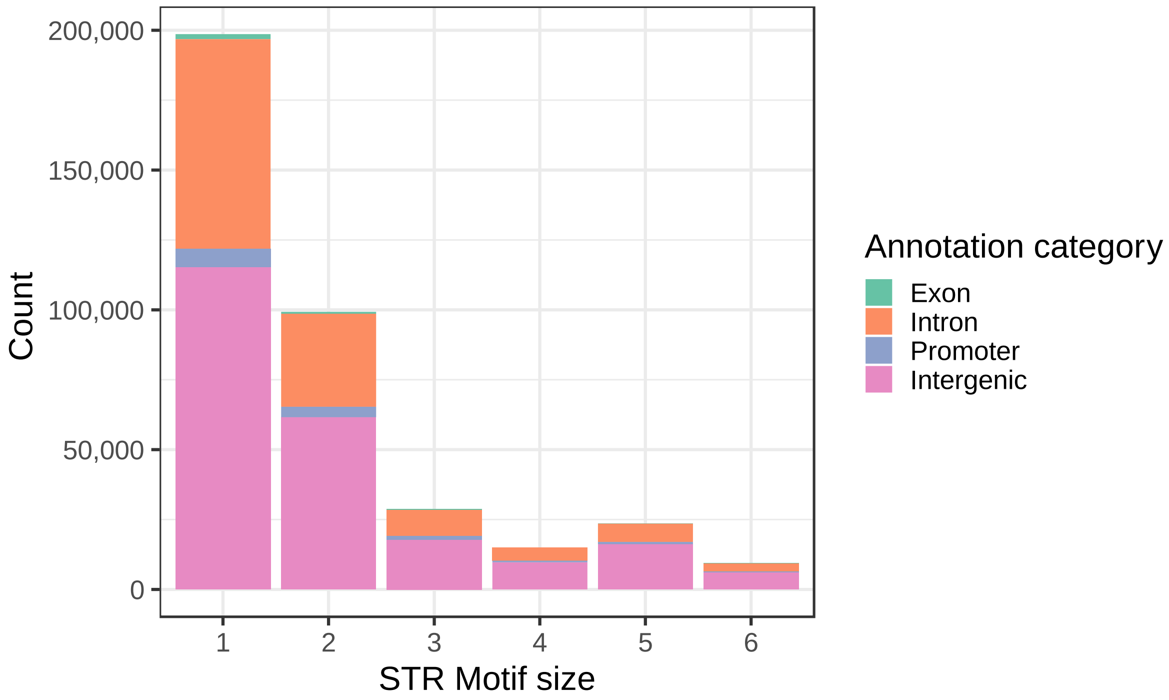


**Figure S7** Number of genotyped STR overlapping four annotation categories. Colours represent different annotation categories.

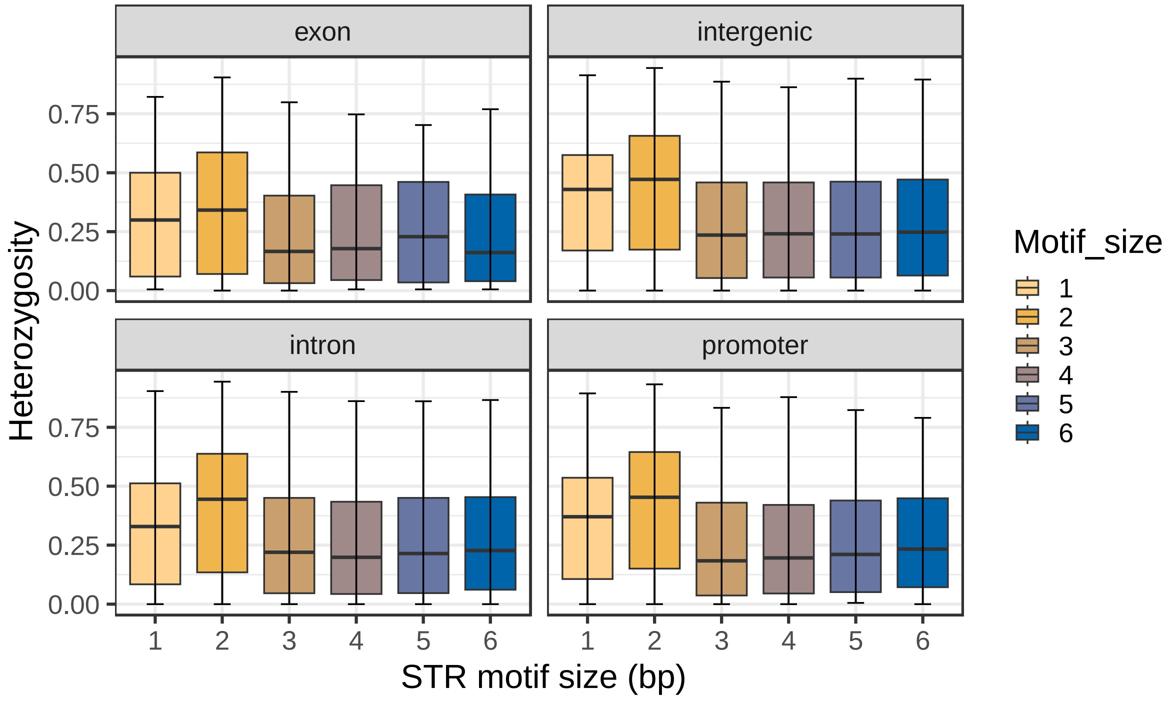


**Figure S8** Heterozygosity of genotyped STR per motif size in each annotation category. Colours indicate different motif sizes.

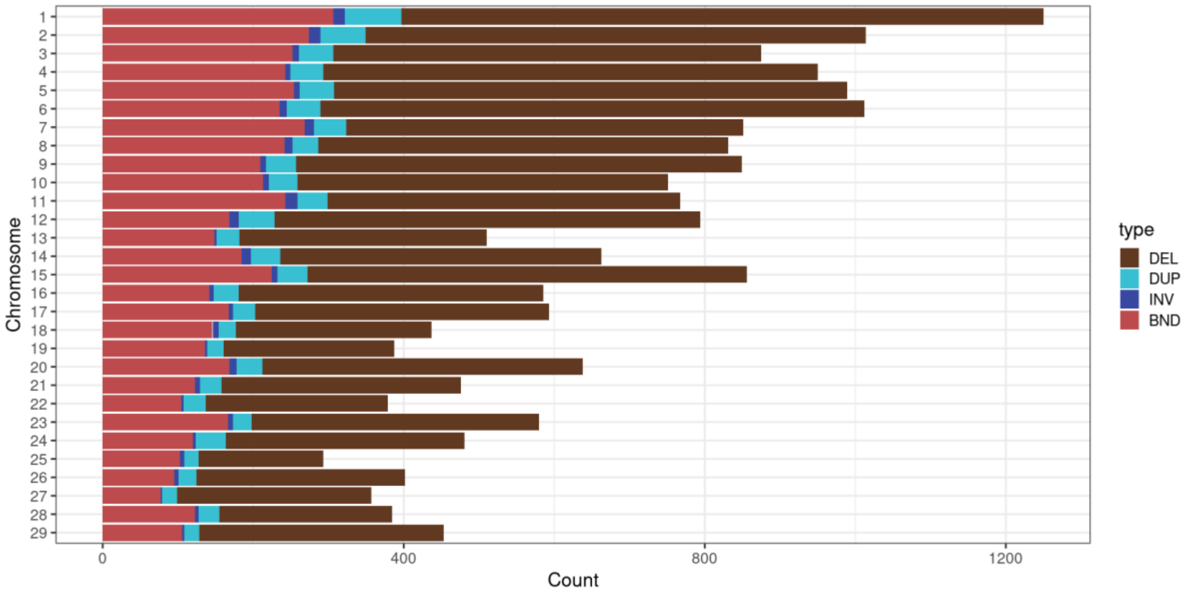


**Figure S9** Cumulative count of SVs found across the 29 autosomes. Colours indicate different SV types.

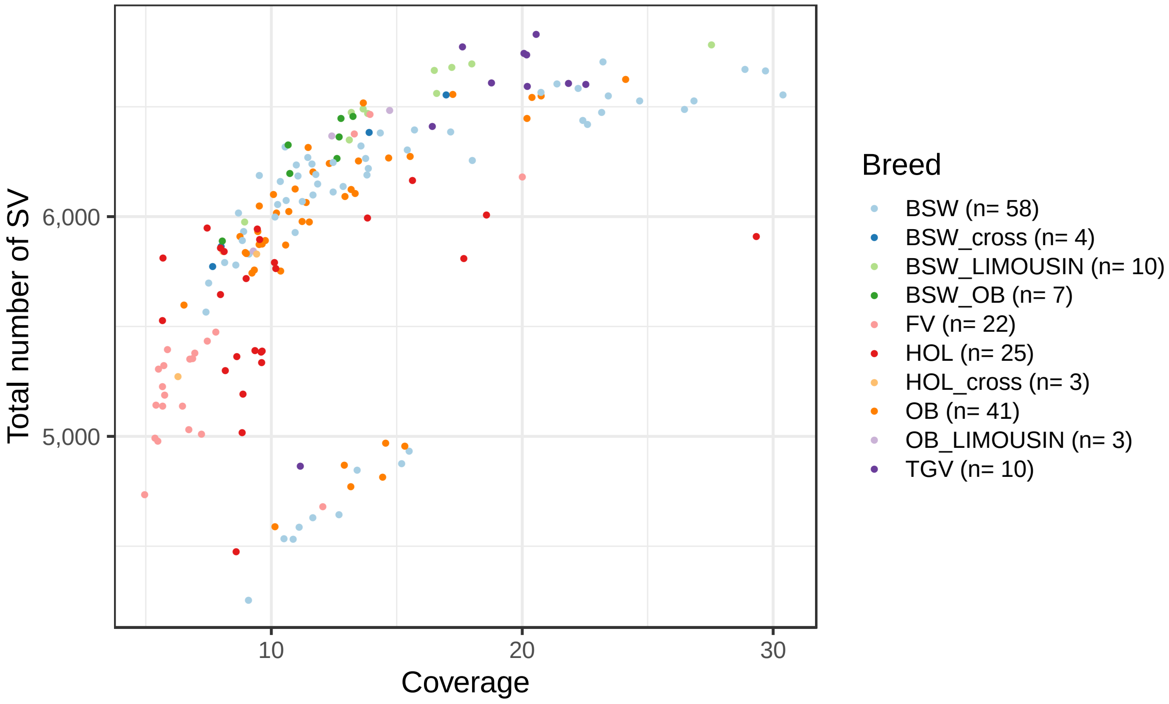


**Figure S10** Number of SVs and autosomal coverage per animal. Colours represent different breeds.

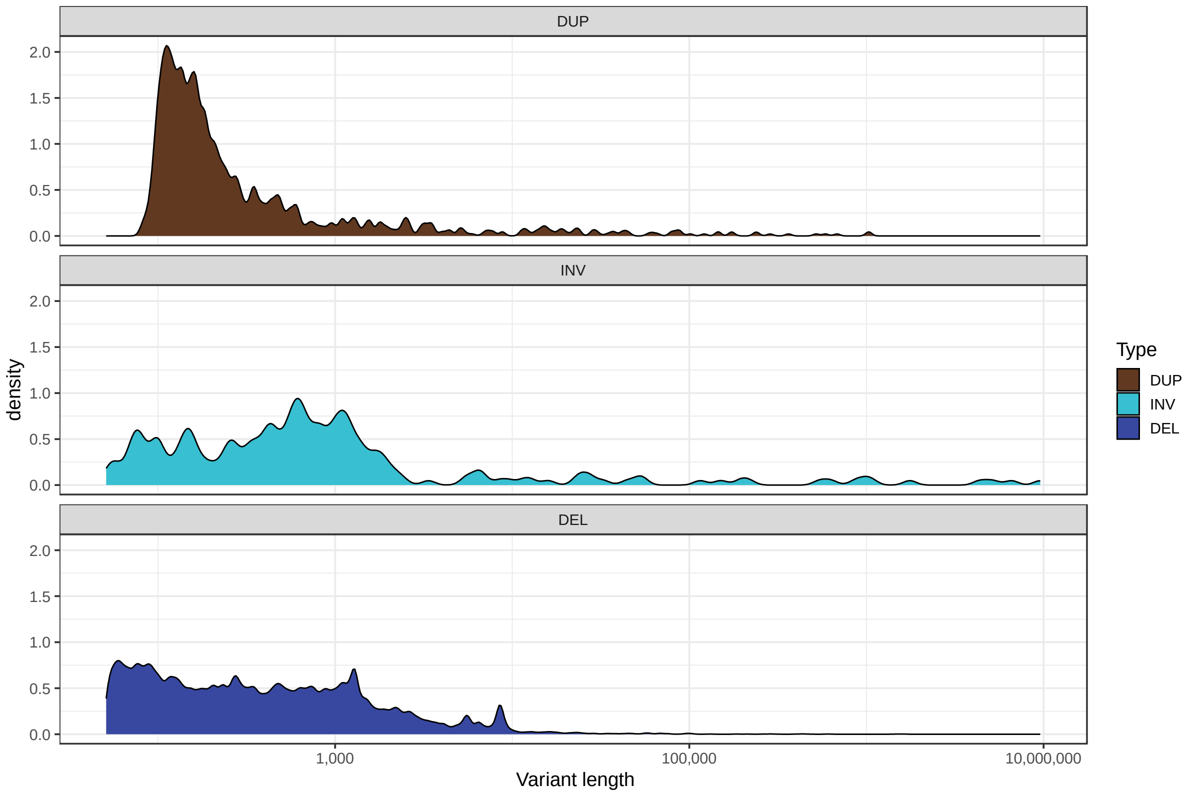


**Figure S11** Density of SVs based on length. BND are not plotted as they all are 1 bp length. Colours indicate different SV types.

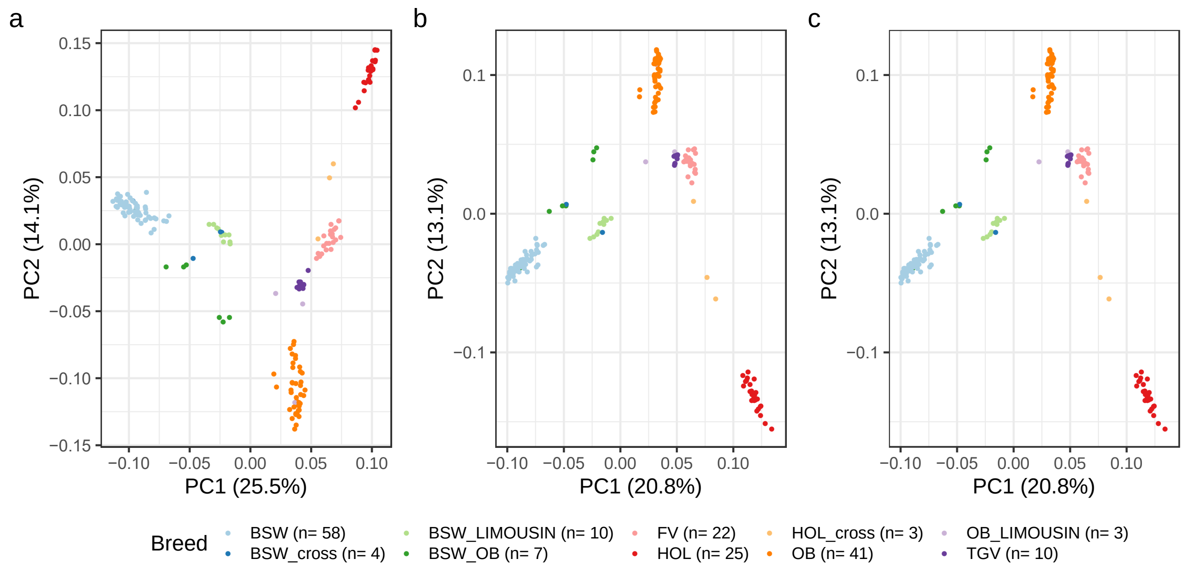


**Figure S12 Principal components analysis.** Visualisation of the top two principal components calculated from a genomic relationship matrix built with (a) SNP, (b) STR, or (c) SV genotypes. Colours represent different breeds.

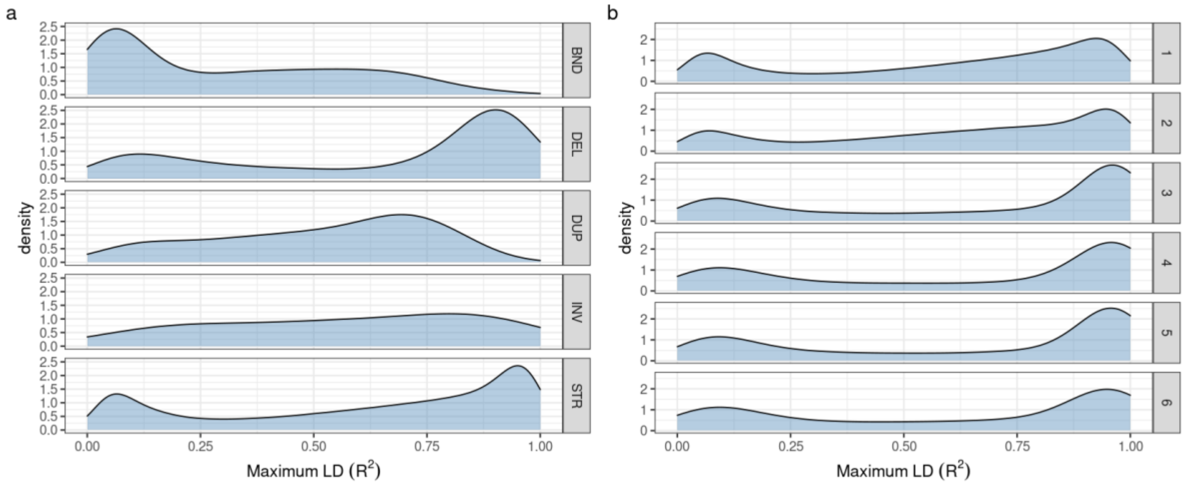


**Figure S13** Distribution of maximum linkage disequilibrium (LD) between SV/STR and SNPs or Indels within 100 kb of each variant. (a) LD between SNPs/Indels and different SV types and STR. (b) LD between SNPs/Indels and different STR motif size.

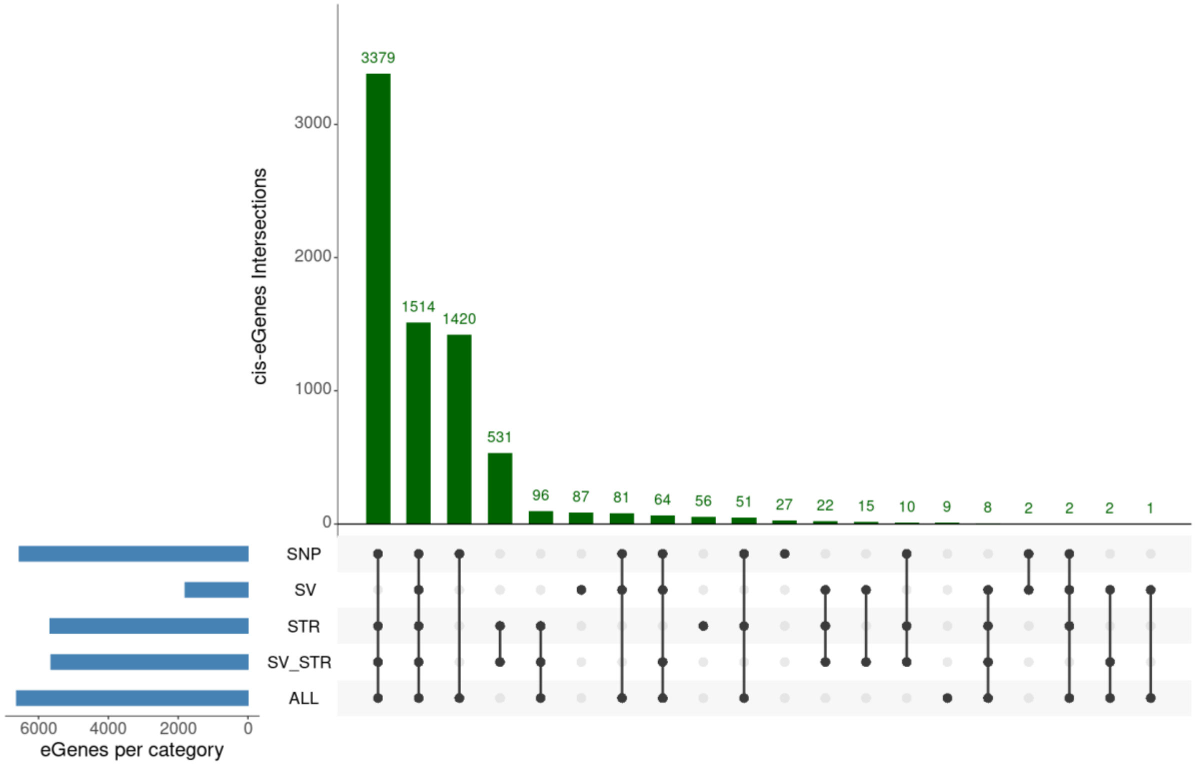


**Figure S14** Overlap of significant cis-eGenes discovered from the ALL, SV-STR and each variant type-specific eQTL analysis.

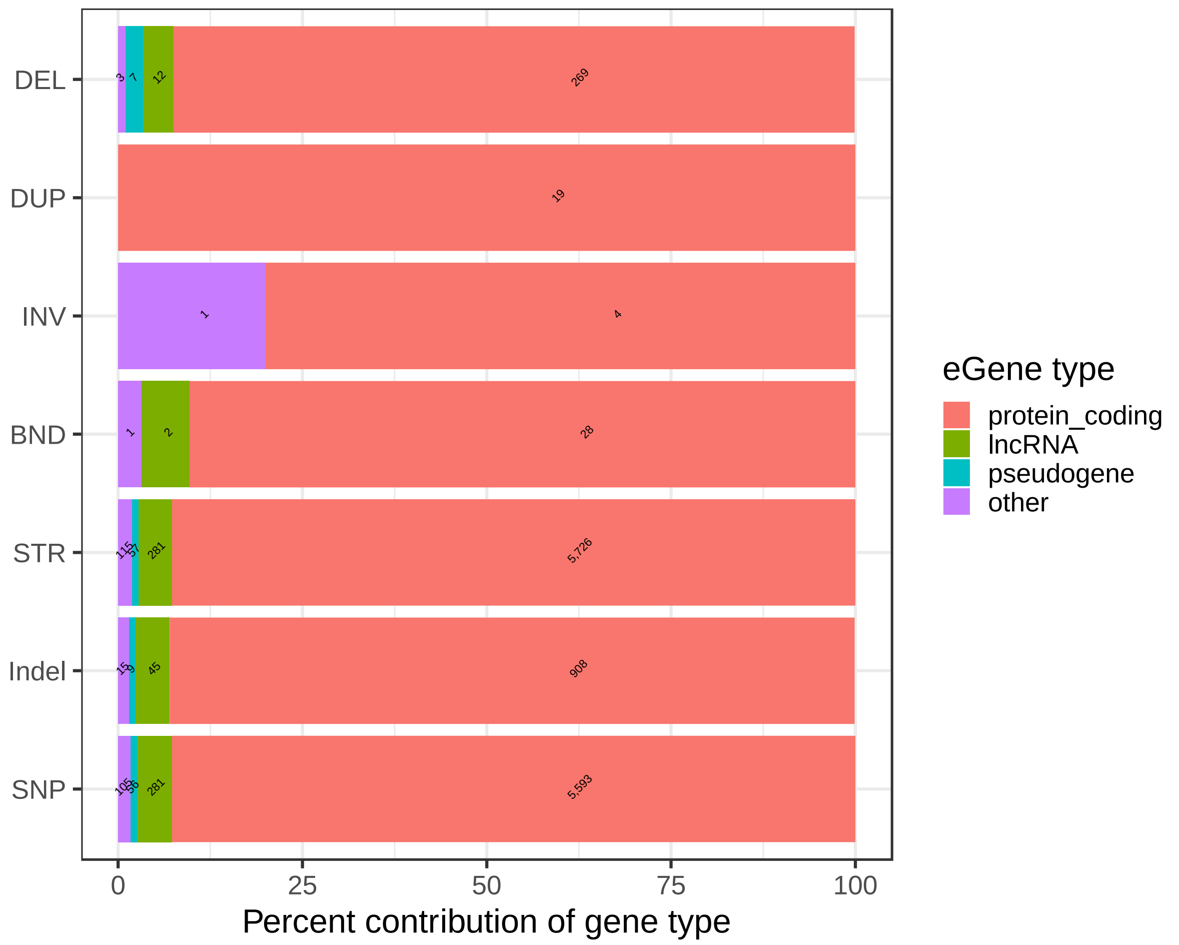


**Figure S15** Fraction of eGenes type associated with respective variant type from the SV-STR eQTL. The numbers inside stacked plot represent the total eQTLs (expression quantitative trait loci) per variant type within each eGene category.

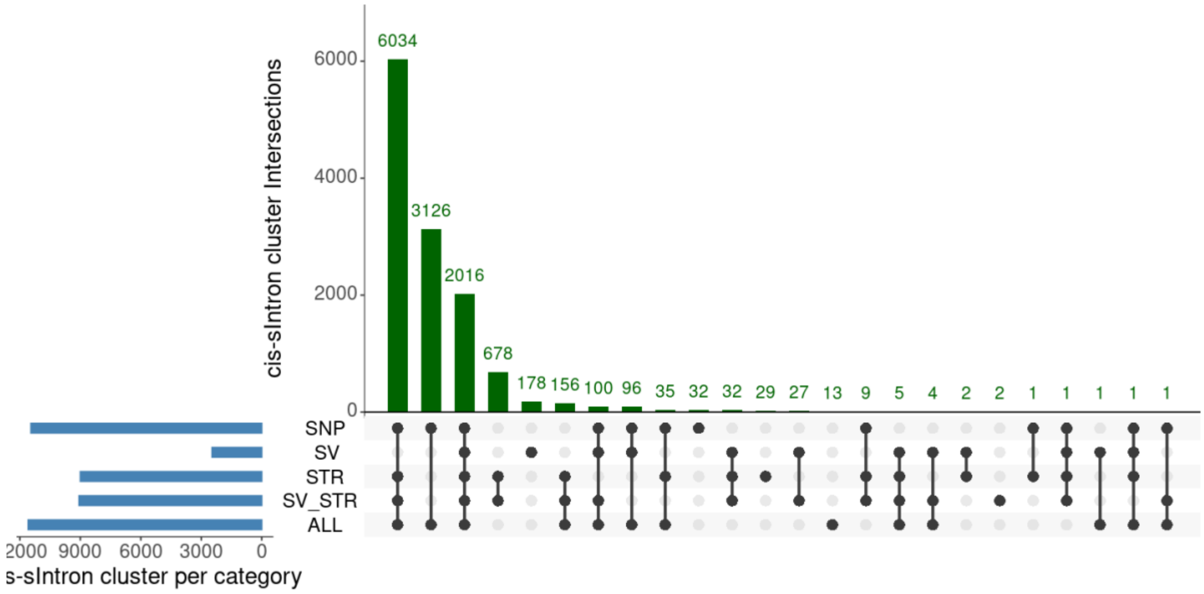


**Figure S16** Overlap of significant sIntron-cluster discovered from each sQTL analysis.

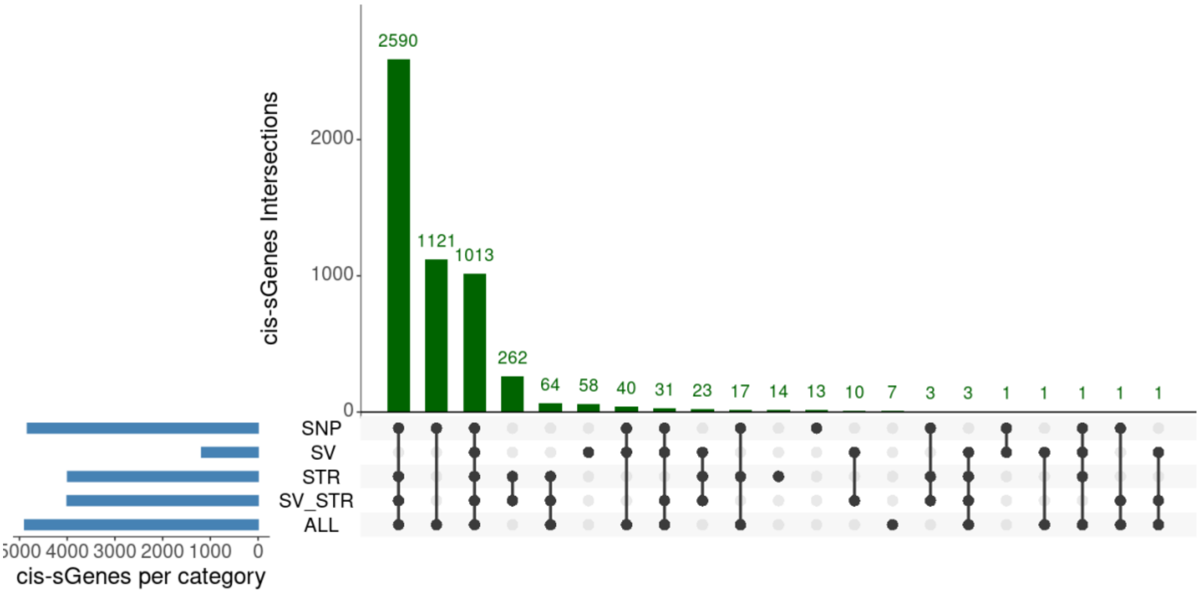


**Figure S17** Overlap of significant cis-sGenes from each sQTL analysis.

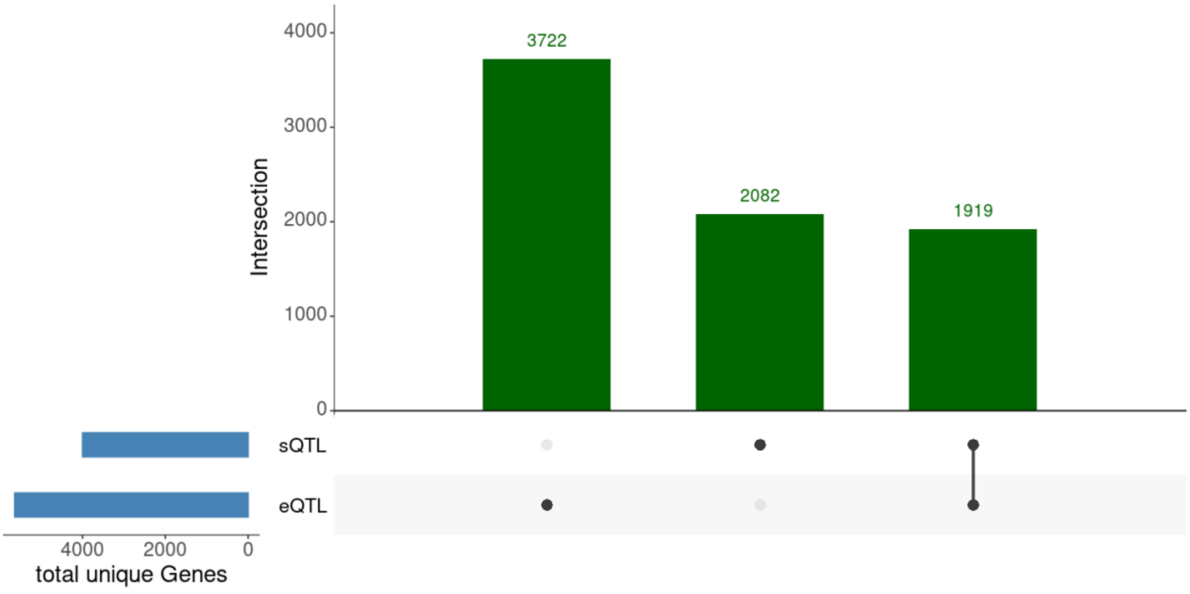


**Figure S18** Overlap of eGenes and sGenes from the SV-STR analyses.

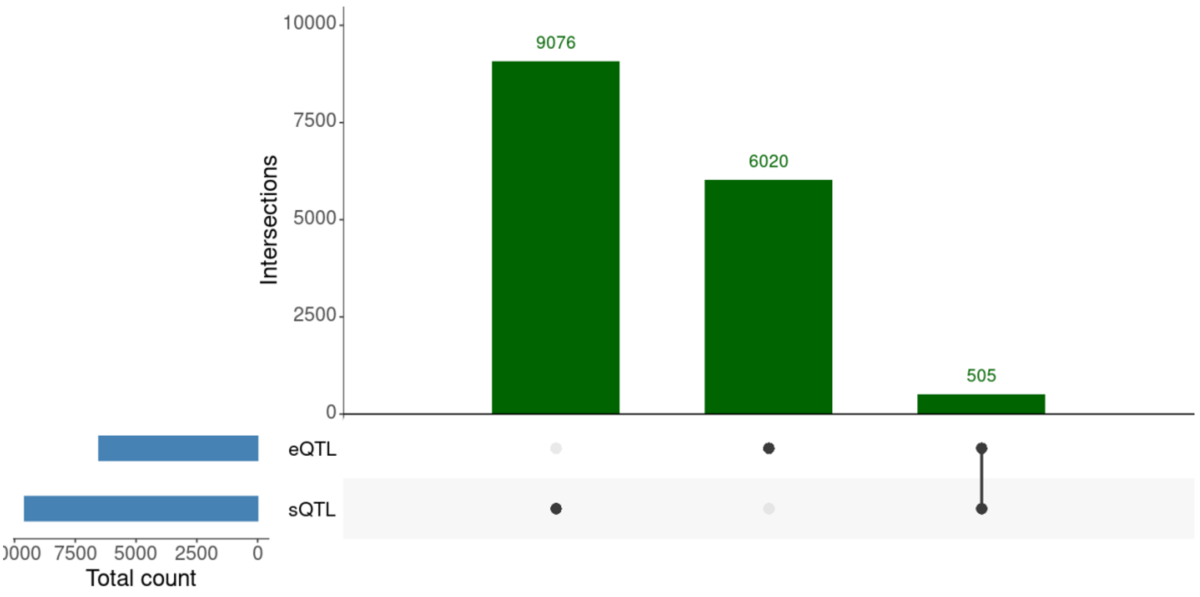


**Figure S19** Overlap of QTL (variant-gene pair) between the eQTL and sQTL SV-STR analyses.
